# A set of *Arabidopsis* genes involved in the accommodation of the downy mildew pathogen *Hyaloperonospora arabidopsidis*

**DOI:** 10.1101/286872

**Authors:** Martina Katharina Ried, Aline Banhara, Fang-Yu Hwu, Andreas Binder, Andrea A. Gust, Caroline Höfle, Ralph Hückelhoven, Thorsten Nürnberger, Martin Parniske

## Abstract

The intracellular accommodation structures formed by plant cells to host arbuscular mycorrhiza fungi and biotrophic hyphal pathogens are cytologically similar but it remains unclear whether these interactions build on an overlapping genetic framework. In legumes, the malectin-like domain leucine-rich repeat receptor kinase SYMRK, the cation channel POLLUX and members of the nuclear pore NUP107-160 subcomplex are essential for symbiotic signal transduction and arbuscular mycorrhiza development. Here we identified members of these three groups in *Arabidopsis thaliana* and explored their impact on the interaction with the oomycete downy mildew pathogen *Hyaloperonospora arabidopsidis (Hpa).* We report that mutations in the corresponding genes reduced the reproductive success of *Hpa* as determined by sporangiophore and spore counts. We discovered that a developmental transition of haustorial shape occurred significantly earlier and at higher frequency in the mutants. Analysis of the multiplication of extracellular bacterial pathogens, *Hpa*-induced cell death or callose accumulation, as well as *Hpa*- or flg22-induced defence marker gene expression, did not reveal any traces of constitutive or exacerbated defence responses. These findings point towards an overlap between the plant genetic toolboxes involved in the interaction with biotrophic intracellular hyphal symbionts and pathogens in terms of the gene families involved.

**Author Summary:** Our work reveals genetic commonalities between biotrophic intracellular interactions with symbiotic and pathogenic hyphal microbes. The majority of land plants engages in arbuscular mycorrhiza (AM) symbiosis with phosphate-acquiring arbuscular mycorrhizal fungi to avoid phosphate starvation. Nutrient exchange in this interaction occurs via arbuscules, tree-shaped fungal structures, hosted within plant root cells. A series of plant genes including the Symbiosis Receptor-like kinase (SYMRK), members of the NUP107-160 subcomplex and nuclear envelope localised cation channels are required for a signalling process leading to the development of AM. The model plant *Arabidopsis thaliana* lost the ability to form AM. Although the ortholog of SYMRK was deleted during evolution, members of the malectin-like domain leucine-rich repeat receptor kinases (MLD-LRR-RKs) gene family, components of the NUP107-160 subcomplex, and an ortholog of the nuclear envelope-localized cation channel POLLUX, are still present in the *Arabidopsis* genome, and *Arabidopsis* leaf cells retained the ability to accommodate haustoria, presumed feeding structures of the obligate biotrophic downy mildew pathogen *Hyaloperonospora arabidopsidis*. We discovered that both of these plant-microbe interactions utilize a corresponding set of genes including the ortholog of POLLUX, members of the NUP107-160 subcomplex and members of the MLD-LRR-RK gene family, thus revealing similarities in the plant program for the intracellular accommodation of biotrophic organisms in symbiosis and disease.

## Introduction

Most land plant species feed carbon sources to arbuscular mycorrhiza (AM) fungi, which in turn deliver phosphate and other nutrients via finely branched intracellular structures called arbuscules [1]. The accommodation of fungal symbionts inside living plant cells involves substantial developmental reprogramming of the host plant cell, initiated by the formation of the pre-penetration apparatus, a transvacuolar cytoplasmic bridge that guides the fungal invasion path [2,3]. The oomycete downy mildew pathogen *Hyaloperonospora arabidopsidis* (*Hpa*) develops intracellular feeding organs, so called haustoria, which exhibit certain structural similarities to the arbuscules of AM fungi [4]. Like arbuscules, haustoria are accommodated inside plant cells and are entirely surrounded by a plant-derived membrane, which keeps the pathogen physically outside the host cytoplasm [5–10]. Structural and functional similarities between accommodation structures for intracellular microbes raised the hypothesis that the corresponding symbiotic and pathogenic associations rely on a similar or even overlapping genetic program [4]. This would imply that filamentous hyphal pathogens exploit an Achilles heel, the presence of the symbiotic program in most land plant species, for their own parasitic lifestyle [11].

In the present study, we tested this hypothesis by focussing on an ancient genetic program comprising the common symbiosis genes (CSGs) [12], which is conserved among angiosperms for the intracellular accommodation of AM fungi [13,14]. In legumes and actinorhizal plants, the CSGs are also required for root nodule symbiosis (RNS) with nitrogen-fixing rhizobia or actinobacteria, respectively [15–17]. In the legume *Lotus japonicus*, the products of some of the CSGs including the MLD-LRR-RK Symbiosis Receptor-like Kinase SYMRK [16,18–20] as well as the nuclear-envelope localised cation channel POLLUX [21,22], are implicated in a signal transduction pathway leading from the perception of microbial signalling molecules at the plasma membrane to the induction of calcium oscillations in and around the nucleus (calcium-spiking), which finally results in the transcriptional activation of symbiosis-related genes and a developmental reprogramming of the respective host cells [1,23]. Furthermore, the nucleoporins NUP85, NUP133 and SEC13 homolog (SEH1) of the NUP107-160 subcomplex are important for the proper establishment of RNS and AM symbiosis [24–27]. The NUP107-160 complex has been implicated in the transport of membrane proteins to the inner membrane of the nuclear envelope [28]. It has been hypothesised that mutations in the NUP107-160 complex may cause a reduced cation channel (e.g. POLLUX) and calcium channel concentration on the inner envelope, which is in accordance with the lack of nuclear calcium spiking in the *NUP107-160* mutants (Jean-Michel Ané, personal communication).

*Arabidopsis thaliana* belongs to the Brassicaceae, a plant lineage that lost specific CSGs and with this the ability to establish AM symbiosis after the divergence of the Brassicales [29–31]. *A. thaliana* can, however, establish a compatible interaction with the hyphal pathogen *Hpa.* The striking similarities between arbuscules and haustoria led us to investigate whether the interaction of *A. thaliana* with *Hpa*, the fungal powdery mildew pathogen *Erysiphe cruciferarum*, which forms intracellular haustoria in *A. thaliana* epidermal leaf cells, or the extracellular bacterial pathogen *Pseudomonas syringae* rely on a similar gene set as the interaction of plants with beneficial hyphal microbes.

## Results

### *A. thaliana* SNUPO mutants reduce the reproductive success of *Hpa*

We inspected the *A. thaliana* genome for the presence of genes related to those required for the interaction with beneficial microbes. Several of the orthologs of key genes required for AM, for example *SYMRK*, *CCaMK* and *CYCLOPS*, are specifically deleted from the *A. thaliana* genome [14,29–32], and thus inaccessible for analysis in the context of the *A. thaliana* interaction with *Hpa*. However, we identified the ortholog of nuclear-localised cation channel *POLLUX* and members of the NUP107-160 subcomplex (S1 + S2 Fig.; Table S1). Furthermore, we identified candidate genes belonging to the MLD-LRR-RK family as close relatives of *LjSYMRK* [16], which we consequently named *SYMRK-homologous Receptor-like Kinase* 1 (*ShRK1*) and *ShRK2* (S1 Fig., S2 Fig., S3 Fig., S4 Fig.; Table S1).

Lines carrying insertional mutant alleles of ***<u>S</u>****hRKs*, selected members of the **<u>NU</u>**P107-160 subcomplex, and the *Arabidopsis* ortholog of ***<u>PO</u>****LLUX* (SNUPO genes) were analysed for their phenotype in the interaction with *Hpa*. In *pollux*, *shrk1*, *shrk2*, the *shrk1* x *shrk2* double mutant, as well as a reference line with a T-DNA insertion in *Phytosulfokine Receptor 1* (*pskr1)* [33,34] or a reference line with an EMS-induced mutation in *Constitutive expression of PR genes 5* (*cpr5*) [35], the reproductive success of *Hpa* isolate NoCo2 - measured as sporangiophore number per cotyledon or spore production - was significantly reduced compared to wild-type plants (Fig. 1 a + c; S5 Fig.).

**Fig. 1:**
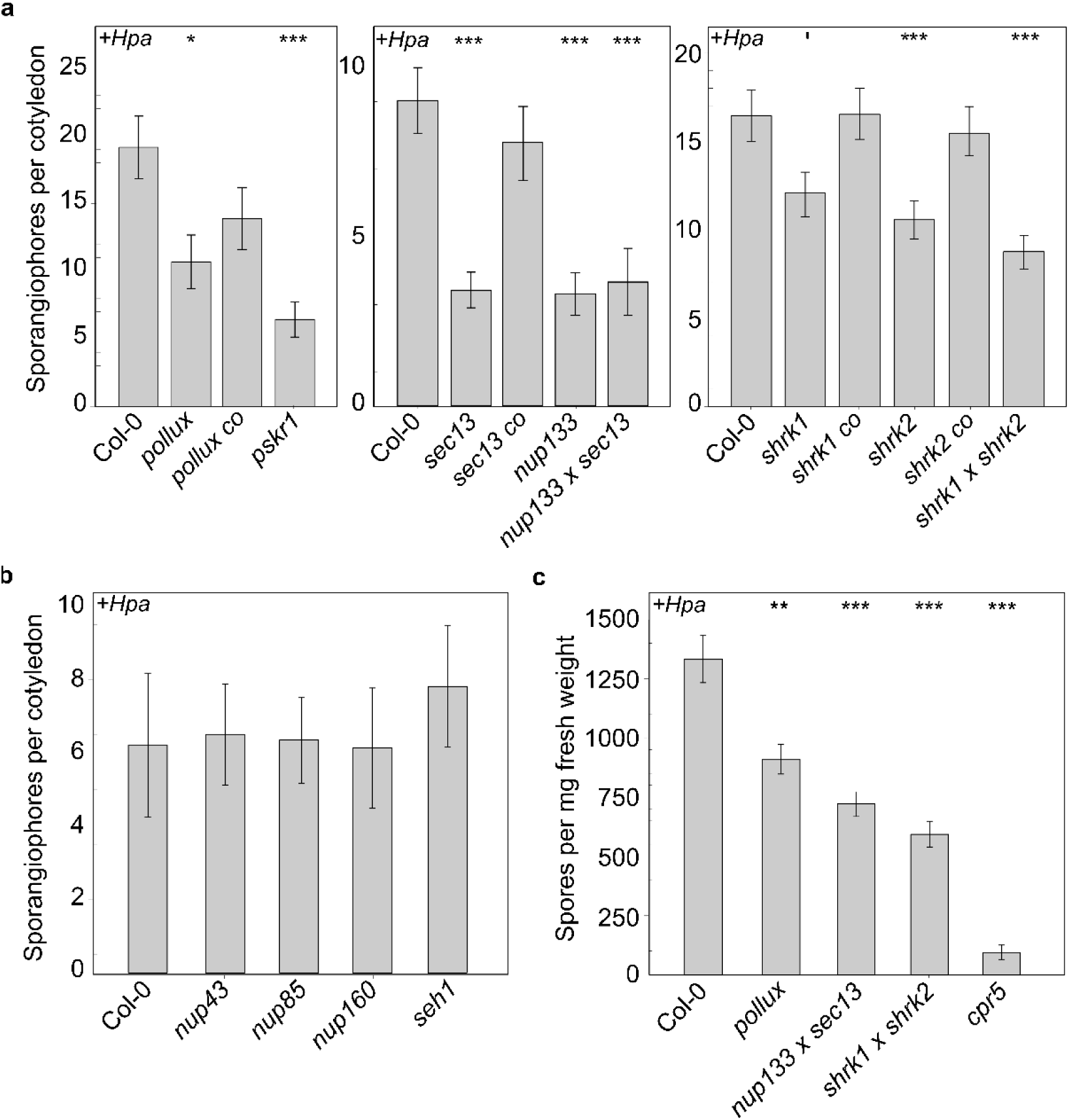
Mutations in *A. thaliana* SNUPO genes reduce the reproductive success of the oomycete downy mildew pathogen *Hpa*. **(a + b)** Bar charts represent the mean number of sporangiophores ± s.e.m on infected cotyledons of *A. thaliana* wild-type (Col-0) or the indicated mutants 4 dpi with *Hpa* isolate NoCo2. n = 14 - 63 **(c)** Bar charts represent the mean number of spores ± s.e.m per mg fresh weight isolated from *A. thaliana* wild-type (Col-0) or the indicated mutants 4 dpi with *Hpa* isolate NoCo2. n = 9. Stars indicate significant differences to Col-0 (Wilcoxon-Mann-Whitney test with Bonferroni-Holm correction; ‘, p = 0.066 *, p < 0.05; **, p < 0.01; ***, p < 0.001). Each experiment was performed at least three times independently.

We analysed a range of NUP107-160 subcomplex members and found that the *sec13* and *nup133* single mutants and the *sec13* x *nup133* double mutant impaired *Hpa* reproductive success (Fig. 1 a + c; S5 Fig.), while lines with T-DNA insertions in *nup43*, *nup85*, *nup160* and *seh1* did not (Fig. 1 b).

All SNUPO genes analysed are constitutively expressed in *Arabidopsis* leaves, in epidermal and mesophyll cells with no consistent clear up- or downregulation in the context of *Hpa* infection [36,37]. The reduced *Hpa* reproductive success could not be explained by a decreased frequency of haustoria formation measured as percentage of host cells contacted by hyphae that harbour an haustorium. Other than a slight decrease in *pollux*, the frequency in the other mutants was indistinguishable from the wild-type (S6 Fig.).

### *A. thaliana* SNUPO mutants display a significantly higher frequency of multilobed *Hpa* haustoria

However, all *A. thaliana* SNUPO mutants, but not *nup43*, *nup85*, *nup160* and *seh1* mutants, exhibited a significant shift in the ratio of haustoria morphologies (Fig. 2; S7 Fig.). At 5 days post infection (dpi) the majority of haustoria in leaves of the wild-type had a globular, single-lobed appearance. Deviations from such morphology, typically characterised by multiple lobes on a single haustorium (“multilobed”), were observed as well. The amount of multilobed haustoria in the SNUPO mutants was significantly increased, a phenomenon that was alleviated in the tested complementation lines (Fig. 2 a + b). In contrast to the SNUPO mutants, the disease resistant *pskr1* mutant did not show any signs of altered haustoria development (Fig. 2 a). In both, wild-type and *shrk1* x *shrk2*, the percentage of multilobed haustoria increased over time, but was significantly higher in *shrk1* x *shrk2* at each analysed time point (Fig. 2 d). To test whether multilobed haustoria are the result of the harsh chemical clearance of the leaf and trypan blue staining procedure, we visualised the haustoria shape *in vivo*. We made use of a stable transgenic *A. thaliana* line expressing RPW8.2-YFP [38] (Fig. 2) and could confirm that multilobed haustoria appeared already in living plant cells (Fig. 2).

**Fig. 2:**
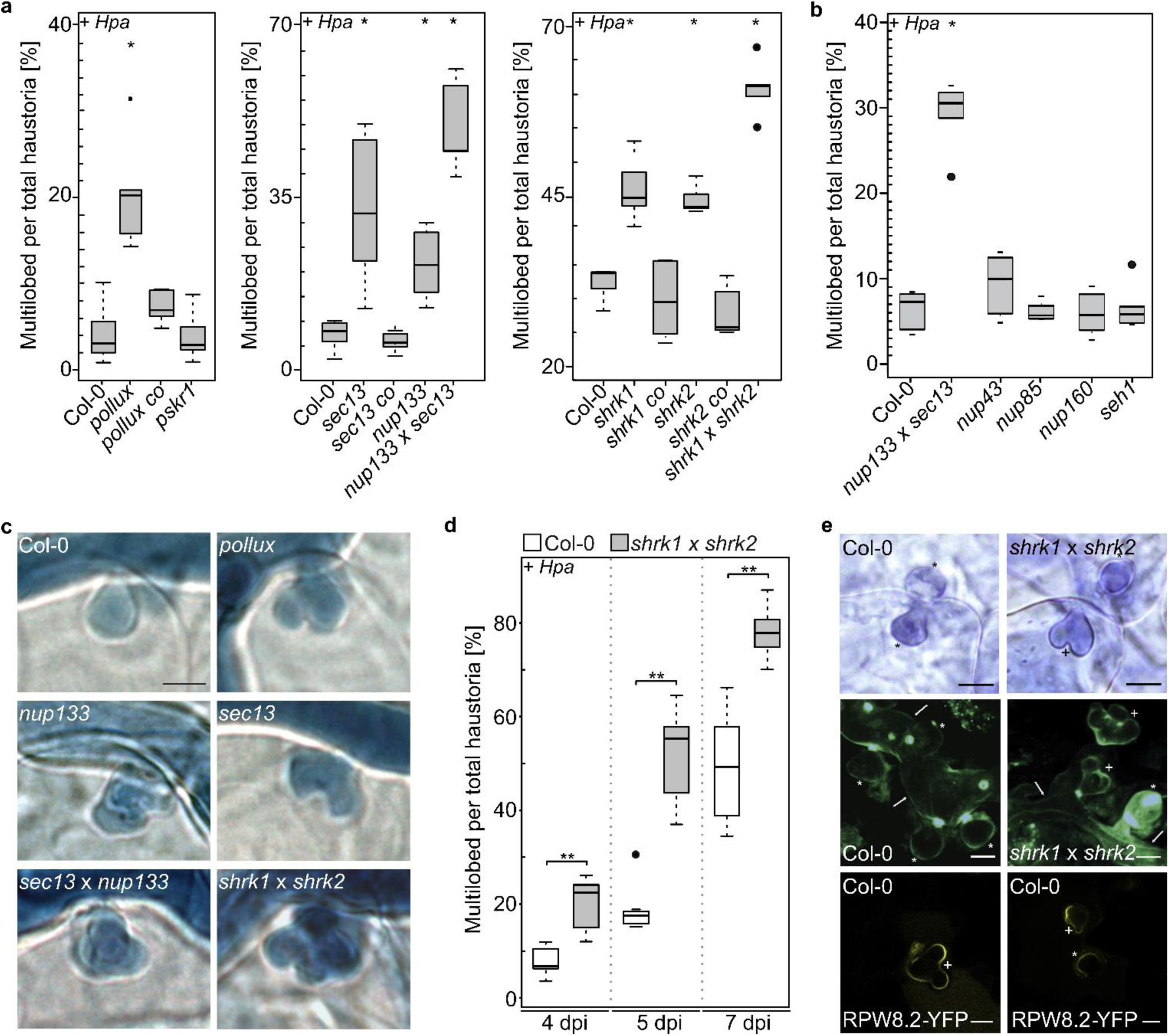
The morphology of *Hpa* haustoria is altered in *A. thaliana* SNUPO mutants. Boxplots represent the percentage of multilobed haustoria among total haustoria on *A. thaliana* wild-type (Col-0), the indicated mutants and transgenic homozygous T3 complementation lines (*pollux co, pPOLLUX:POLLUX; sec13 co, pSEC13:SEC13; shrk1 co, pUbi:ShRK1; shrk2 co, pUbi:ShRK2; LjSYMRK, pUbi:Lotus japonicus SYMRK-mOrange*) 5 dpi with *Hpa* isolate NoCo2 **(a + b)**, or 4, 5 and 7 dpi with *Hpa* isolate NoCo2 **(d)**. For each genotype, at least ten independent stretches of hyphae per leaf have been analysed on at least 5 leaves. On average, 1100 haustoria have been analysed per genotype. **(c)** Representative pictures of intracellular haustoria are displayed. Bars = 25 μm. **(e)** Upper panel: Light microscopy of Technovit section of *A. thaliana* wild-type (Col-0) or the *shrk1 x shrk2* double mutant 7 dpi with *Hpa* isolate NoCo2. Bars, 10 μm. Middle panel: Confocal light scanning microscopy of leaves of *A. thaliana* wild-type (Col-0) or the *shrk1 x shrk2* double mutant stained with aniline-blue 5 dpi with *Hpa* isolate NoCo2. For every haustorium, the entry point can be identified by the formation of the usually bright callose neck. Bars, 25 μm. Lower panel: Confocal light scanning microscopy of leaves of *A. thaliana* wild-type (Col-0) expressing RPW8.2-YFP, which localises to the perihaustorial membrane 7 dpi with *Hpa* isolate NoCo2. *, round haustoria; +, multilobed haustoria; arrows, hyphae. Bars, 10 μm. Black circles, data points outside 1.5 IQR of the upper/lower quartile; bold black line, median; box, Interquartile Range (IQR); whiskers, lowest/highest data point within 1.5 IQR of the lower/upper quartile. Stars indicate significant differences to Col-0 (Wilcoxon-Mann-Whitney test with Bonferroni-Holm correction; *, p < 0.05; **, p < 0.01; ***, p < 0.001). Each experiment was performed at least three times independently.

### *ios1* mutants of ecotypes Ler and Col-0 differ in the frequency of multilobed haustoria

*L. japonicus* SYMRK belongs to the LRR I-RK family of receptor kinases, which consists of 50 members in *A. thaliana* [39,40]. Of these, *Impaired Oomycete Susceptibility 1* (*IOS1*) has been implicated in defence-related signalling [40,41]. On the one hand, *IOS1* supports the infection of an obligate biotrophic oomycete pathogen, but on the other hand, is important for the resistance against bacteria [40,41]. Interestingly, we could only observe enhanced frequency of multilobed haustoria in an *ios1* mutant of the *A. thaliana* ecotype Ler but not in an *ios1* mutant of Col-0, which might imply that different paralogs are important for plant-microbe interaction in different ecotypes (S8 Fig.).

### The reproductive success of a powdery mildew pathogen appears unaltered on *A. thaliana* SNUPO mutant leaves

To investigate the specificity of the role of SNUPO genes in plant-pathogen interactions, we infected SNUPO mutants with the haustoria-forming powdery mildew fungus *E. cruciferarum*. We did not observe a consistent reduction in the reproductive success of *E. cruciferarum* (Fig. 3; S9 Fig.). A morphological comparison of haustoria shape was not attempted because of the much more complex and highly variable haustoria morphology in this interaction.

**Fig. 3:**
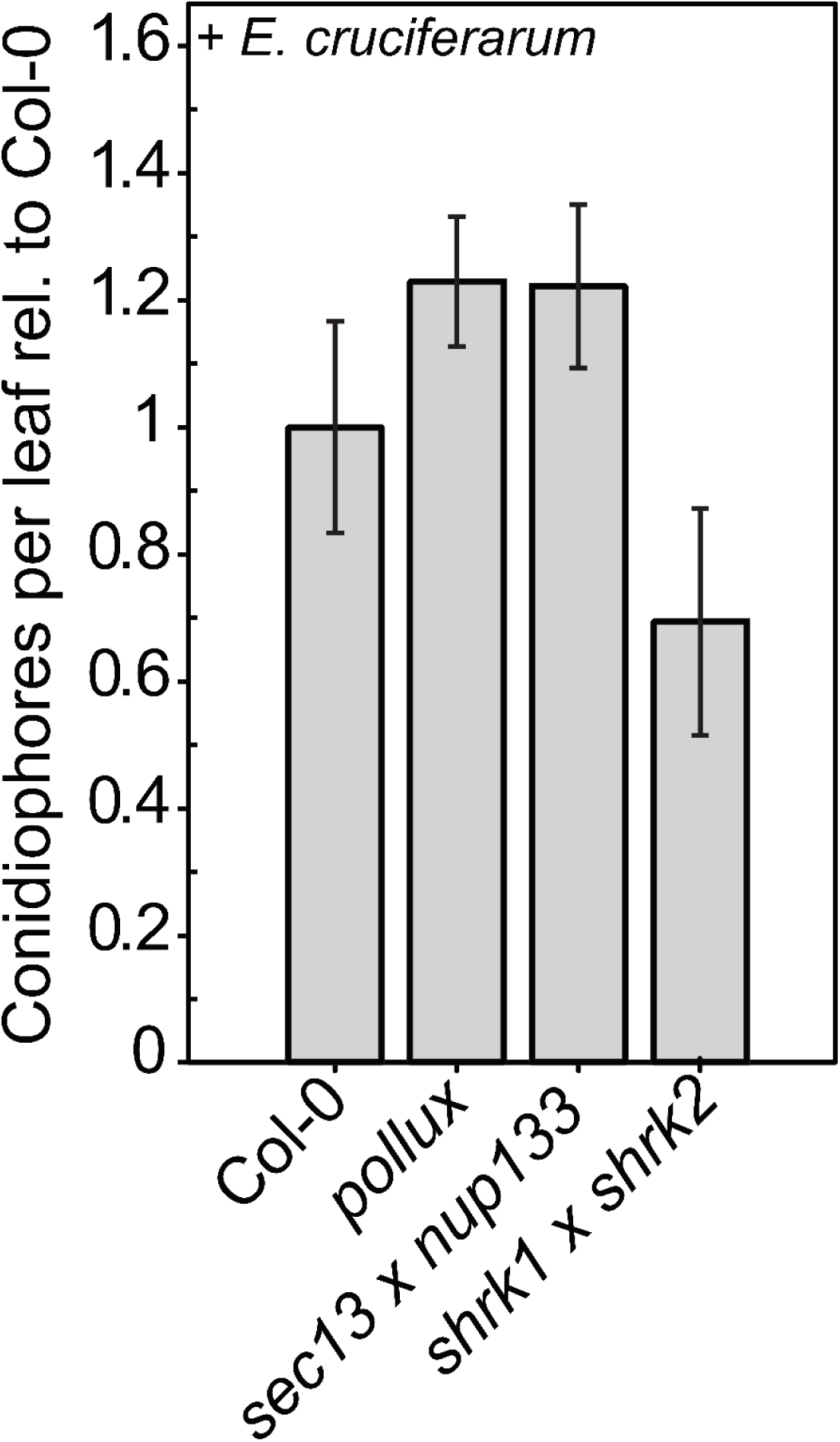
Mutations in *A. thaliana* SNUPO genes do not impair the reproductive success of the fungal powdery mildew pathogen *E. cruciferarum*. Bar charts represent the mean number of conidiophores ± s.e.m on leaves of the indicated mutants relative to the wild-type (Col-0) 5 dpi with *E. cruciferarum*. At least ten colonies per leaf on at least 6 leaves have been counted. No significant differences to Col-0 were detected (Wilcoxon-Mann-Whitney test with Bonferroni-Holm correction; *, p < 0.05; **, p < 0.01; ***, p < 0.001).

### *A. thaliana* SNUPO mutants do not exhibit constitutive or exacerbated activation of defence responses

Plant evolution resulted in several defence strategies against microbial pathogens. For example, PAMP triggered immunity (PTI) is initiated by the perception of pathogen-associated molecular patterns (PAMPs) or plant-derived damage-associated molecular patterns (DAMPs) and results in plant defence responses accompanied by the activation of defence marker genes or callose deposition [42]. To analyse whether the increased *Hpa* resistance of the SNUPO mutants is a result of deregulated defence responses, we investigated the basal transcript levels of the three PAMP-induced marker genes - *Pathogenesis-Related gene 1* (*PR1*), a marker gene for salicylic acid (SA)-mediated resistance [43], *Ethylene Response Factor 1* (*ERF1*), a marker gene for ethylene-mediated resistance [44] and the plant defensin *PDF1.2a*, a marker gene for jasmonic acid (JA)- and ethylene-mediated resistance [45] in the absence of pathogenic attack. Transcript levels in the tested mutants under non-challenged conditions did not differ from transcript levels in the wild-type (Fig. 4 a, black circles). To examine whether SNUPO mutants show enhanced activation of defence marker genes, we analysed the transcript levels of *PR1*, *ERF1* and *PDF1.2a* in plants infected with *Hpa.* All tested genes were upregulated to a similar extent in the SNUPO mutants and the wild-type (Fig. 4 a, empty circles). Only *PDF1.2a* transcript levels were significantly lower in *Hpa-*infected *shrk1 x shrk2* plants than in infected wild-type plants, however, it is unlikely that this is the reason for the increased pathogen resistance of the double mutant. Moreover, the ability of *Hpa* to suppress callose deposition around the haustorial neck region [46] was not disturbed in the SNUPO mutants compared to the wild-type (Fig. 4 b).

**Fig. 4:**
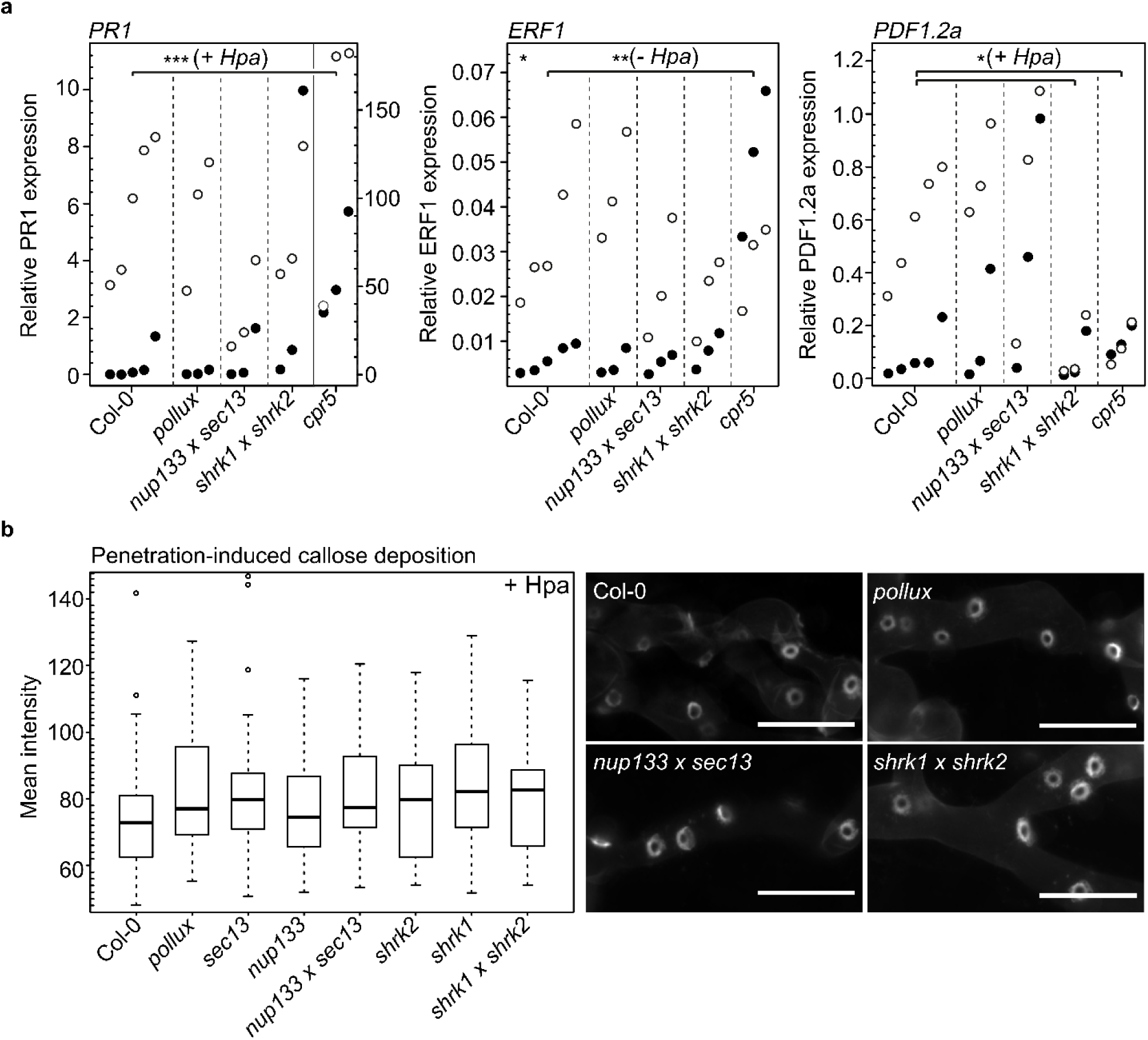
Activation of defence responses in response to *Hpa* infection is unaltered in *A. thaliana* SNUPO mutants. **(a)** Expression of *PR1, ERF1* or *PDF1.2a* relative to the housekeeping genes *TIP41-like* and *PP2A* in mock-treated samples (black circles) or in *Hpa*-infected samples (empty circles) was determined in three biological replicates for each genotype by qRT-PCR. Stars indicate significant differences to Col-0 (Dunnett’s Test with Bonferroni correction; *, p < 0.05). **(b)** Boxplots represent the mean intensity of callose deposition in leaves of *A. thaliana* wild-type (Col-0) or the indicated mutants 4 dpi with *Hpa* isolate NoCo2. n = 27 - 72. Circles, data points outside 1.5 IQR of the upper/lower quartile; bold black line, median; box, IQR; whiskers, lowest/highest data point within 1.5 IQR of the lower/upper quartile. No significant differences to Col-0 were detected (Wilcoxon-Mann-Whitney test with Bonferroni-Holm correction; *, p < 0.05; **, p < 0.01; ***, p < 0.001). Representative pictures of haustoria-associated callose deposition at the neck band of *Hpa* haustoria growing on the indicated mutants are shown on the right. Bars, 25 μm.

In addition to the expression levels of defence marker genes, we investigated other symptoms typically associated with deregulated immune responses. We could not observe any developmental or growth defects, which are typically a result of the hyper-activation of the SA-dependent defence pathway [35], in SNUPO mutants grown side-by-side with the wild-type and the dwarf mutant *suppressor of npr1-1, constitutive 1* (*snc1* [47]) (S10 Fig.). Furthermore, we analysed constitutive and *Hpa*-induced cell death responses in SNUPO mutants and the wild-type (Fig. 5). Most of the non-infected leaves did not display any sign of cell death (Fig. 5, first column), but dark-blue stained dead cells were sporadically observed with no significant differences between non-infected leaves of both wild-type and SNUPO mutants (Fig. 5, second column and upper graph). In infected leaves of the SNUPO mutants, cell death is detected in, or adjacent to, haustoria-containing cells in a frequency indistinguishable from or even lower than the wild-type (Fig. 5, third column and lower graph). In all genotypes, infected leaves contain hyphal strands growing in the absence of any cell death (Fig. 5 fourth column).

**Fig. 5:**
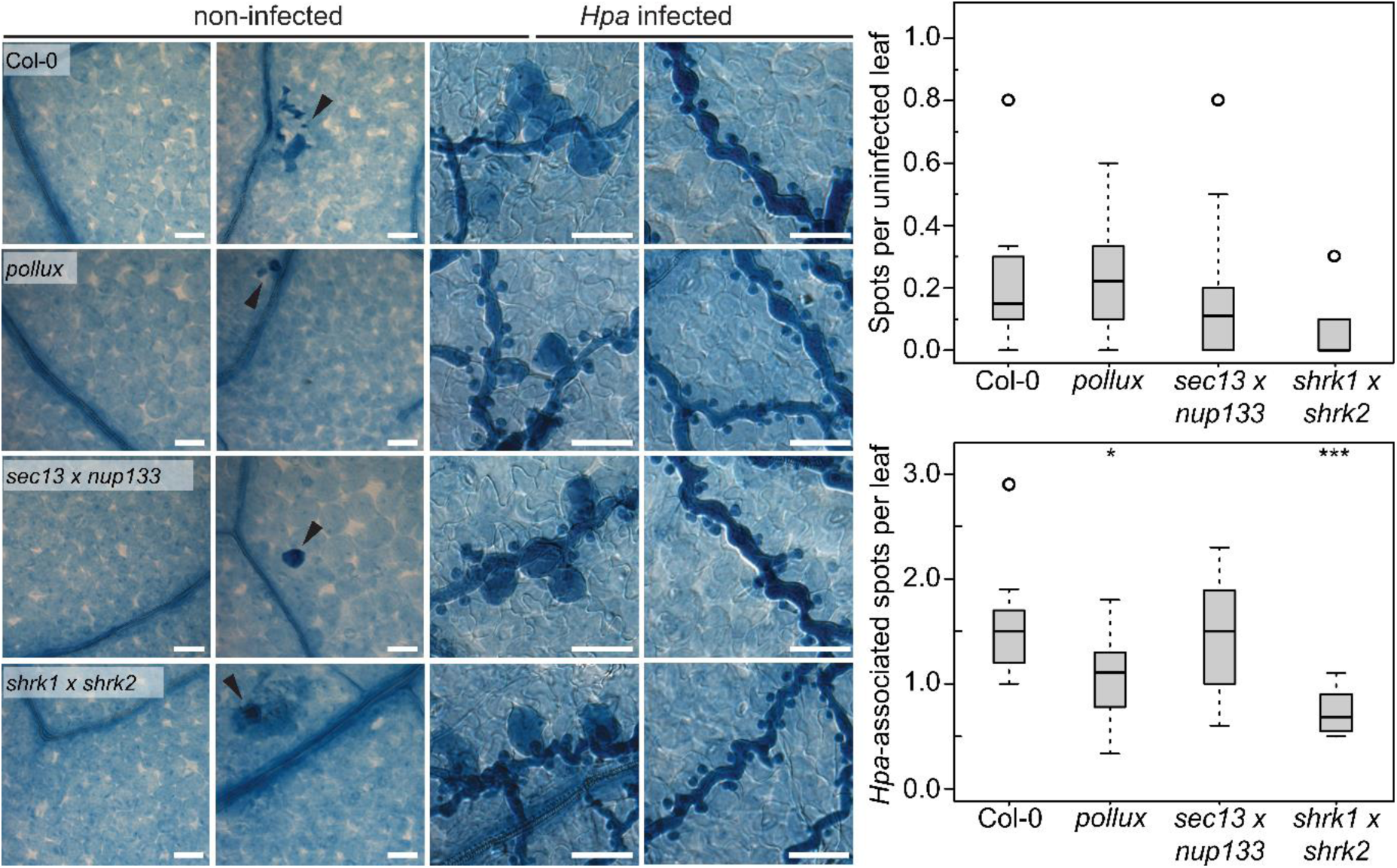
Cell death responses are unaltered in *A. thaliana* SNUPO mutants. Differential interference contrast microscopy of *A. thaliana* wild-type (Col-0) and CSG mutant leaves stained with trypan-blue lactophenol 4 dpi with *Hpa* isolate NoCo2. First column, non-infected without cell death; second column, non-infected with random cell death; third column, infected with cell death in, or adjacent to, haustoria-containing cells; fourth column, infected without cell death. Bars, 25 μm. Boxplots represent the number of random (upper graph) or *Hpa*-associated (lower graph) cell death spots per leaf on *A. thaliana* wild-type (Col-0) or the indicated mutants 5 dpi with *Hpa* isolate NoCo2. On each leaf four to ten spots have been analysed and a total of five leaves per genotype were investigated. Circles, data points outside 1.5 IQR of the upper/lower quartile; bold black line, median; box, IQR; whiskers, lowest/highest data point within 1.5 IQR of the lower/upper quartile. Stars indicate significant differences to Col-0 (Wilcoxon-Mann-Whitney test with Bonferroni-Holm correction; *, p < 0.05; **, p < 0.01; ***, p < 0.001).

### flg22-induced gene expression and resistance to *P. syringae* is unaltered in *A. thaliana* SNUPO mutants

To examine whether mutation of SNUPO genes has an effect on signalling related to PAMP-triggered immunity, we inspected the transcript levels of *Flg22-induced Receptor-like Kinase 1* (*FRK1* [48]) and *ERF1* in the wild-type, the SNUPO mutants and a *fls2* mutant (Fig. 6) in response to the bacterial flagellin-derived peptide flg22 [49]. Transcript levels in mock-treated samples were not different in the wild-type and the analysed mutants, except for the *shrk1* mutant that displayed a slight increase in basal *FRK1* transcript levels, which was not detectable in the *shrk1 x shrk2* double mutant, and the *nup133 x sec13* double mutants that contained slightly decreased basal levels of *ERF1* transcripts. The deviations observed for individual mutants are unlikely responsible for the increased *Hpa* resistance. Six hours after flg22 treatment the tested genes were all upregulated to the same extent in the SNUPO mutants and the wild-type (Fig. 6 a). Furthermore, we investigated the growth behaviour of *P. syringae* on the wild-type and on SNUPO mutants (Fig. 6 b). *P. syringae* pv. *tomato* (*Pto*) DC3000 induces the activation of SA-dependent defence signalling in the host, and deregulation of this pathways impairs *P. syringae* resistance [50]. The growth of *Pto* DC3000 wild-type or the avirulent *ΔAvrPto/PtoB* strain was unaltered on the *A. thaliana* SNUPO mutants, providing further evidence that they do not exhibit constitutive or enhanced activation of SA-dependent defences (Fig. 6 b).

**Fig. 6:**
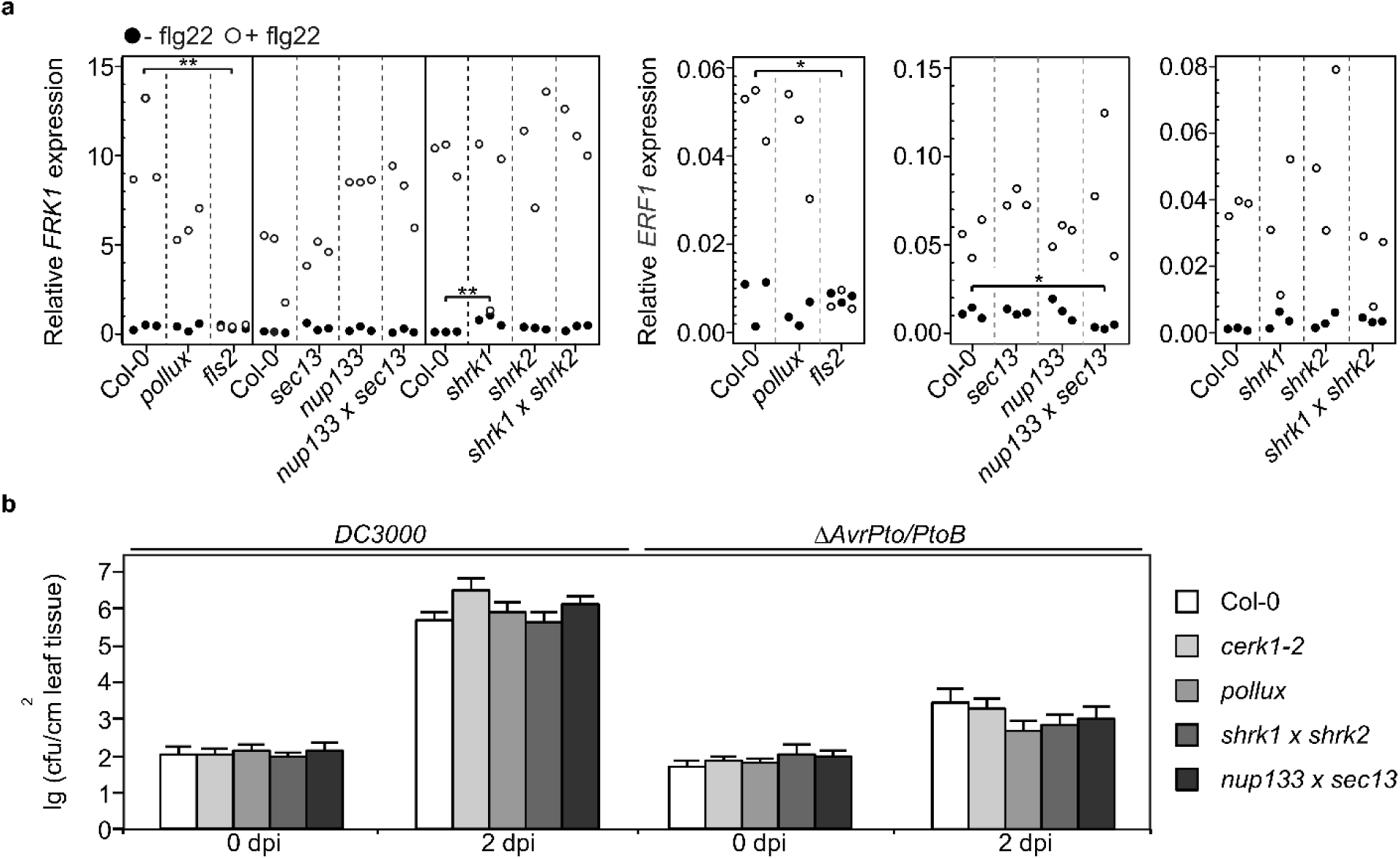
Expression of flg22-inducible genes as well as growth of the bacterial pathogen *Pseudomonas syringae* are not affected in *A. thaliana* SNUPO mutants. (a) Expression of *FRK1* and *ERF1* relative to the housekeeping genes *TIP41-like* and *PP2A* in mock-treated samples (black circles) or in flg22-treated samples (empty circles) was determined in three biological replicates for each genotype by qRT-PCR. The *fls2* mutant served as negative control. Stars indicate significant differences to Col-0 (Dunnett’s Test with Bonferroni correction; *, p < 0.01; **, p < 0.001). **(b)** Bacterial density of *Pto* DC3000 or *Pto* DC3000 Δ*AvrPto/PtoB* on *A. thaliana* wild-type (Col-0) or the indicated mutants was determined as colony forming units per cm^2^ 2 dpi. Data represent means ± s.d. of six replicate measurements/genotype/data point. Results from one representative of at least four independent experiments are shown.

## Discussion

Obligate biotrophic hyphal pathogens like *Hpa* that form haustoria inside living plant cells depend on the vitality of their host cell for their own survival and reproduction. The plant genes and developmental programs of the host that are supporting the formation and maintenance of oomycetal haustoria are largely unexplored. Here we tested the hypothesis that the biotrophic pathogen *Hpa* utilises a gene set related to that enabling intracellular accommodation in the context of AM symbiosis.

### SNUPO mutants reveal a link between the timing of haustorial shape shifts and reproductive success

We observed - based on sporangiophore and spore counts - that *A. thaliana* SNUPO mutants are impaired in supporting the reproduction of the oomycete pathogen *Hpa*. This was associated with a shift in the haustoria morphology. Intriguingly, we observed congruent phenotypes in mutants of diverse protein classes with the only connector between these protein classes being their reported involvement in root endosymbioses. The frequency of multilobed haustoria increased in the wild-type as well as in the mutants over time revealing a clear connection between haustoria morphology and the age of the interaction, which is in line with a previously observed continuous growth of *Albugo* and *Hpa* haustoria over time (Eric Kemen and Marco Thines; independent personal communication). As the frequency of multilobed haustoria was significantly higher in the *A. thaliana* SNUPO mutants at all time points analysed, it appears that haustoria growth is accelerated in the SNUPO mutants. It has been postulated that haustoria are the main avenue for nutrient acquisition from the host [51]. Considering the agricultural impact of biotrophic hyphal pathogens, surprisingly little is known about the precise function of haustoria and changes thereof during haustoria development. Senescence of haustoria has been associated with encasement which was discussed to likely reduce their functionality [52]. Therefore, it is possible that the accelerated formation of multilobed haustoria is a sign of senescence and thus directly responsible for the reduction in *Hpa* reproductive success. This scenario would suggest a role of *Arabidopsis* SNUPO genes in maintaining the single lobed, and presumably functional, stage of *Hpa* haustoria. In an opposite scenario, the surface increase from single lobed to multilobed haustoria may benefit the oomycete and the genes under study are involved in delaying the progress of haustoria development into the multilobed stage. However, this scenario is not compatible with the reduced sporangiophore and spore count on the mutants.

### SNUPO mutants exhibit no traces of altered plant defence responses or regulation

Based on the analysis of a wide range of defence symptoms in the classical PTI assays and three different pathosystems, we conclude that *A. thaliana* SNUPO mutants display unaltered levels and frequencies of defence responses. Consequentially, the impairment of the *Hpa* interaction is not due to deregulated defence in these mutants.

### *Arabidopsis* SNUPO mutants are specifically impaired in the *Hpa* interaction

We were unable to detect differences in the interaction of *A. thaliana* SNUPO mutants with the haustoria-forming powdery mildew fungus *E. cruciferarum* in comparison to wild-type plants. This may be due to the different cell types targeted by *Hpa* and *E. cruciferarum* for haustoria formation. *Hpa* initially colonizes epidermal cells and then progresses to the mesophyll [53], in which the vast majority of haustoria are formed, while *E. cruciferarum* forms haustoria solely within epidermal cells [54]. In addition, the genetic requirements for the intracellular accommodation of fungal and oomycete pathogens may differ. The *A. thaliana Mildew resistance Locus O* (*MLO)* gene, for instance, is an epidermal compatibility factor required for powdery mildew fungus penetration [55], with no role in the *Hpa* interaction reported to date. However, it should be noted that the haustoria of *E. cruciferarum* are structurally very complex and irregularly shaped. Because of this inherent polymorphy, it is possible that we missed subtle structural changes caused by the SNUPO mutations.

### Genetic overlaps between symbiotic and pathogen biotrophic interactions

In contrast to the vast knowledge on genes contributing to disease resistance, relatively few genes have been identified that facilitate pathogen colonization on *Arabidopsis*, such as *PMR4*, *PMR5*, *PMR6* [56–58], *DMRs* [59,60], *MYB3R4* [36], and *IOS1* [40]. While *dmr3* and *dmr6* mutants also exhibited aberrantly shaped haustoria, these mutants strongly differ from the consistent phenotype of the SNUPO mutants. In *dmr6*, oomycete growth is arrested after the formation of the first haustoria and the expression of several defence-related genes is elevated in unchallenged *dmr6* mutants [59,60]. *Dmr3* mutants are more resistant to the bacterial pathogen *P. syringae* pv. *tomato*, they exhibit elevated *PR1* expression, and *dmr3* and *dmr6* mutants show increased resistance to the powdery mildew pathogen *Golovinomyces orontii* [59]. Taken together, *dmr3* and *dmr6* mutants confer broad-spectrum resistance and display perturbed defence-related gene expression [59,60]; these data suggest that *DMR3* and *DMR6* – in contrast to *Arabidopsis* SNUPO genes - are rather involved in general plant disease resistance than in compatibility with and the accommodation of a hyphal organism.

It is a long-standing hypothesis that plant pathogens exploit an Achilles heel of the plant, genetic pathways for the intracellular accommodation of mutualistic symbionts such as AM fungi or nitrogen-fixing bacteria [4,11]. Recently the oomycete pathogen *Phytophthora palmivora*, which forms haustoria that only last for a few hours, has emerged as a hemi-biotrophic model system in two independent laboratories to test this hypothesis in legume mutants. This pathogen quickly progresses to a necrotrophic phase and structural alterations in haustoria development are thus less likely to affect pathogen fitness [61–64]. Huisman and colleagues did not detect any alterations in the infection of *Medicago truncatula* mutant roots of the CSGs *DMI1*, *DMI3* and *IPD3* by *P. palmivora* [64]. Considering that the lifetime of infected cells in the *P. palmivora* interaction is reduced, it appears that the transient biotrophic phase of *P. palmivora* is too short and progresses too fast into the necrotrophic phase to allow the haustorial maintenance functions of the CSGs – or a related gene set - to take effect [65,66]. Therefore, the beneficial effect of these genes or their homologs for hyphal pathogens may only be detectable in longer lasting biotrophic relationships. The too short lifetime of the host cell may also explain the absence of a detectable phenotype in *A. thaliana* SNUPO mutants and *L. japonicus* CSG mutants infected with the beneficial fungus *Piriformospora indica* [67]. Colonisation with the endophytic fungus *Colletotrichum tofieldiae*, is controlled by the plant phosphate starvation response system and *C. tofieldiae* only increases plant fertility and promotes plant growth under phosphorus-limiting conditions [68]. It will be an interesting task of future research to investigate whether *Arabidopsis* SNUPO genes are implicated in the interaction with this beneficial microbe. In contrast, for the symbiosis genes for which a mutant phenotype has been described - namely *RAM2* and *RAD1* - the mutation was not affecting haustoria structure [61–63]. Mutants in the glycerol-phosphate acyl-transferase gene *RAM2* of the model legume *M. truncatula*, for example, had a reduced abundance of small cutin monomers and did not elicit the formation of appressoria by *P. palmivora* at the surface of roots and caused an altered infection behaviour of the AM fungus within the root cortex [63]. This observation also implies that a compatibility factor can be exploited by microbial pathogens and, likewise, by beneficial symbionts. Similarly, mutants of the GRAS protein gene *RAD1*, share a role in AM symbiosis and the *P. palmivora* interaction [61].

Our data revealed genetic commonalities of symbiosis and disease in the formation of intracellular accommodation structures at a later developmental stage of the plant-microbe association. A key difference between our study and the previous ones that failed to observe a mutant phenotype in pathogenic interactions is the longer lasting biotrophic phase in the *A. thaliana* - *Hpa* interaction. Although strongly suggested by the function of the CSGs, it remains unclear whether *A. thaliana* SNUPO genes are similarly involved in a signal transduction pathway directly supporting oomycete development. It will be therefore interesting to identify the mechanistic commonalities between symbiotic and pathogenic interactions with hyphal organisms that are controlled by corresponding gene sets.

### Retention of SNUPO genes in the *Arabidopsis* genome

The loss of AM symbiosis in *A. thaliana* and in four other independent plant lineages was correlated with the absence of more than 100 genes with potential roles in AM [29–31,69]. While the exploitation of symbiotic programs by pathogens might explain the consistent deletion of CSGs from five independent plant lineages, it raises the question, which evolutionary forces retained SNUPO genes in the *Arabidopsis* genome. A housekeeping function was not revealed since no pleiotropic developmental phenotypes were observed in the mutants. While, it might be possible that these genes limit *Hpa* colonisation, thereby forcing the oomycete to sporulate earlier or more profusely, our results leave us with the unexpected finding that the only detected role for the SNUPO genes in *A. thaliana* is the support of an oomycete. It will be interesting to find out whether ecological conditions exist, under which oomycete colonization might provide a selective advantage to the host plant, or whether SNUPO genes are also involved in the accommodation of beneficial microbes like *C. tofieldiae* justifying their retention in the *Arabidopsis* genome [70].

## Materials and Methods

### Seed sterilization and plant growth

All *A. thaliana* mutants described in the manuscript were of Col-0 ecotype, except for *ios1*, which was either of Col-0 or Ler ecotype. Seeds were obtained from “The Nottingham Arabidopsis Stock Centre” - NASC [71] or the GABI-DUPLO double mutant collection [72]. For *in vitro* experiments, *A. thaliana* seeds were sterilized by incubation in sterilisation solution (70% ethanol, 0.05% Tween20) for 5 min, followed by incubation in 100% ethanol for 2 min. For *Hpa* infection, seeds were directly germinated in soil and grown for two weeks under long day conditions (16 h light, 22°C, 85 μmol m^−2^ s^−1^). For *E. cruciferarum* inoculation, plants were grown in a 2:1 soil/sand mixture. Seeds were stratified for 48 h at 4°C prior to transfer into a growth chamber (10 h light, 120 μmol m^−2^ s^−1^ light, 22°C day, 20°C night, 65% relative humidity). For elicitor treatment, seeds were placed on half-strength MS plates [73], stratified for 48 h at 4°C in the dark, and grown for 8 days under long day conditions (16 h light, 23°C, 85 μmol m^−2^ s^−1^).

### *A. thaliana* stable transformation

Floral dipping of *A. thaliana* was performed as described previously [74].

### Pathogen assays and phenotypic analyses

To collect spores for inoculation, *A. thaliana* wild-type (Col-0 or Ler) leaves with sporulating *Hpa* isolate NoCo2 or *Hpa* isolate Waco9, respectively, were harvested seven days post inoculation (dpi) and placed into 10 ml deionized H_2_O and vortexed for 2 s in 15 ml reaction tubes. The spore suspension was then filtered through a Miracloth filter and sprayed onto 12-days-old plants using a spraying device. Subsequently, plants were placed into trays and covered with wet translucent plastic lids. Trays were sealed to maintain high humidity, and plants were grown under long day conditions (16 h light, 18°C, 85 μmol m^−2^s^−1^).

For spore counting, *A. thaliana* wild-type (Col-0) and mutant seedlings were harvested 5 dpi. Ten seedlings per background were harvested into reaction tubes containing 1 ml deionized H_2_O and vortexed for 3 min. Spores were counted with a Fuchs Rosenthal chamber. For sporangiophore counting and the investigation of haustoria shape and penetration efficiency cotyledons or leaves were harvested and stained in 0.01% trypan-blue-lactophenol for 3 min at 95°C and 5 h at room temperature, followed by overnight clearing in chloral hydrate (2.5 g/ml). Samples were mounted in glycerol for subsequent differential interference contrast microscopy with a Leica DMI6000B. For sporangiophore counting, a minimum of 50 cotyledons per genotype and replicate were analysed and the number of sporangiophores per infected cotyledon was plotted. For investigation of haustoria shape and penetration efficiency, a minimum of five leaves per genotype and replicate were analysed. On each leaf, the percentage of multilobed haustoria per total haustoria or the percentage of haustoria-containing cells per cells contacted by hyphae was calculated for 10 individual strands of hyphae. On average, 1100 haustoria have been analysed per genotype and experiment. The mean for each leaf was calculated and plotted.

*E. cruciferarum* was grown on *A. thaliana* wild-type (Col-0) to maintain aggressiveness and on susceptible *phytoalexin deficient 4* (*pad4*) mutants [50] for elevated conidia production. Plants were placed under a polyamide net (0.2 mm^2^) and inoculated at a density of 3 - 4 conidia mm^−2^, by brushing conidia off of *pad4* mutant leaves through the net. Two leaves per plant were harvested, cleared and kept in acetic acid (25%) until microscopic analysis. Leaves were stained in acetic acid (25%) ink (1:9) (Königsblau, Pelikan, 4001), washed in water, placed in water containing a few drops of Tween20, washed in water again, and analysed under a bright-field microscope. For each replicate, conidiophores per colony were counted on 10 colonies per leaf, on 5 - 10 leaves per genotype.

Bacterial strains *Pto* DC3000 or *Pto* DC3000 *ΔAvrPto/AvrPto* were grown and used for infection assays on leaves of 4 - 5 weeks old *A. thaliana* plants as described previously [75,76].

Each experiment was performed at least three times independently.

### Observation of haustoria

Leaves of *A. thaliana* wild-type (Col-0) or the *shrk1* x *shrk2* double mutant were harvested at 5 dpi, cleared in 10% KOH for 5 min, stained with 0.05% aniline blue in 67 mM K_2_HPO_4_ for 20 min and observed with a Leica SP5 confocal light scanning microscope. Images were edited using ImageJ with the “volume viewer” plugin [77].

For Technovit sections, *A. thaliana* wild-type (Col-0) or mutant leaves were infected with *Hpa* as described above and harvested at 7 dpi. Leaves were fixed with 3.7% formaldehyde and dehydrated by incubating samples in 30%, 50%, 70% and 100% ethanol. Samples were embedded in Technovit 7100 according to the manufacturer’s instruction. A Leica RM2125 rotary microtome was used to cut 7 μm sections. Sections were stained in 0.01% trypan-blue-lactophenol for 3 h at 37°C, followed by clearing in chloral hydrate (2.5g/ml) and subsequent differential interference contrast microscopy with a Leica DMI6000B.

*A. thaliana* wild-type (Col-0) plants expressing *RPW8.2-YFP* [38] were infected with *Hpa* as described above, harvested at 10 dpi and observed with a Leica TCS SP 5 confocal laser scanning microscope equipped with a 63x NA 1.2 water-immersion objective.

### Analysis of oomycete-associated callose deposition

Oomycete-associated callose deposition was analysed on cotyledons of *A. thaliana* wild-type (Col-0) or SNUPO mutants. Leaves were harvested at 4 dpi, cleared in 10% KOH for 5 min, stained with 0.05% aniline blue in 67 mM K_2_HPO_4_ for 20 min and mounted in glycerol for observation with a Leica DMI6000B with CFP filter settings. Regions of interest (ROIs) covering the neck bands were selected in ImageJ [77] and the signal intensity of each neck band was calculated from single ROIs and mean intensities were plotted. n = 27 - 72.

### Elicitor treatment

For pre-incubation, 8 days old seedlings were transferred to a 12-well plate (3 seedlings/well represent one biological replicate) with half-strength liquid MS medium [73] supplemented with 1% sucrose and incubated overnight under long-day conditions (16 h light, 22°C, 100 μmol m^−2^ s^−1^; 8 h dark, 18°C) and 100 rpm shaking. On the following day the medium was exchanged, half of the samples were supplemented with 1 μM flg22, and the other half was kept in half-strength MS to serve as mock controls. Plants were then incubated for 6 h at 22°C and 100 rpm shaking. For every genotype, three biological replicates of treated and non-treated samples were harvested and immediately frozen in liquid N_2_ for subsequent RNA extraction.

### RNA Extraction and qRT-PCR analysis

RNA extraction was performed using the Spectrum™ Plant Total RNA kit (Sigma-Aldrich), followed by DNase I treatment (amplification grade DNase I, Invitrogen™) to remove genomic DNA. First strand cDNA synthesis was performed from 250 ng total RNA using the SuperScript® III First-Strand Synthesis SuperMix (Invitrogen™) with oligo(dT) primers. qRT-PCR was performed in 20 μL reactions containing 1x SYBR Green I (Invitrogen™) in a CFX96 Real-time PCR detection system (Bio-Rad). PCR program: 2’-95°C; 40 x (30’’-95°C; 30’’-60°C; 20’’-72°C); melting curve 95°C – 60°C – 95°C. A primer list can be found in S3 Table. Expression levels of target genes were normalized against the housekeeping genes *TIP41-like* and *PP2A* [78]. For every genotype, three biological replicates represented by two technical duplicates each were analysed.

### Gene structure and phylogenetic analyses

Complete annotated genomic sequences were obtained from The *Arabidopsis* Information Resource (TAIR - www.arabidopsis.org) for *A. thaliana*, and from the KDRI website (Kazusa DNA Research Institute, Japan; http://www.kazusa.or.jp/lotus/) and the GenBank for *L. japonicus*. BLAST searches were performed on TAIR (http://arabidopsis.org/Blast/index.jsp) with the *L. japonicus* genomic CSG sequences as query. The protein domain organization and the exon-intron structure of the *A. thaliana* homologs of *POLLUX*, *SEC13*, *NUP133* (GenBank accession number: KM269292), *ShRK1* and *ShRK2* are identical to that of their *L. japonicus* counterparts. By sequencing a PCR product amplified from *A. thaliana* wild-type (Col-0) cDNA, we demonstrated that, contrary to the TAIR prediction, this was also the case for *NUP133*. The curated sequence has been submitted to TAIR. For phylogenetic studies, protein sequences of *A. thaliana* and *L. japonicus* MLD-LRR-RKs were aligned using MAFFT 7 [79] with default settings. Alignments were used to create phylogenetic trees at the CIPRES web-portal with RAxML 8.2.10 [80] for maximum likelihood analyses. For RAxML, the JTT PAM matrix for amino acid substitutions was chosen.

### Statistics and data visualisation

All statistical analyses and data plots have been performed and generated with R version 3.0.2 (2013-09-25) “Frisbee Sailing” [81], using the packages “Hmisc” [82], “car” [83], “multcompView” [84] and “multcomp” [85] or Excel. For statistical analysis, data was either subjected to the nonparametric Wilcoxon-Mann-Whitney test with Bonferroni-Holm correction using Col-0 samples as control group, or was power transformed to improve normality and a one-way ANOVA followed by a Dunnett’s Test with Bonferroni correction was performed using Col-0 samples as control group.

## Acknowledgements

We thank Ruth Eichmann for performing pilot experiments with *Erysiphe cruciferarum*, Birgit Kemmerling for kindly sending us *pskr-1* seeds, Dirk Metzler for help with statistical analyses and Rosa Elena Andrade Aguirre for help with the phylogenetic analyses.

## Supporting Information

**S1 Fig.**
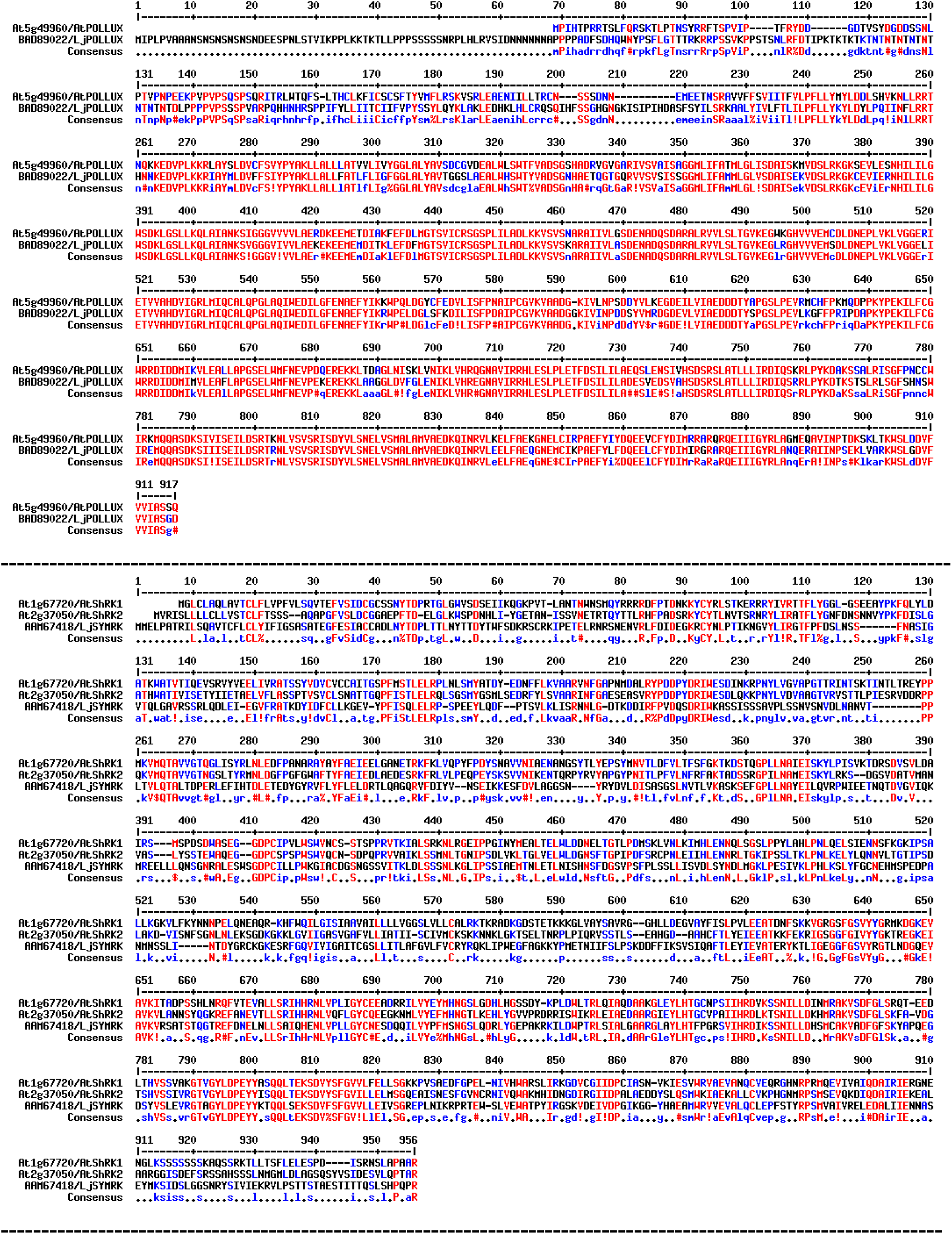
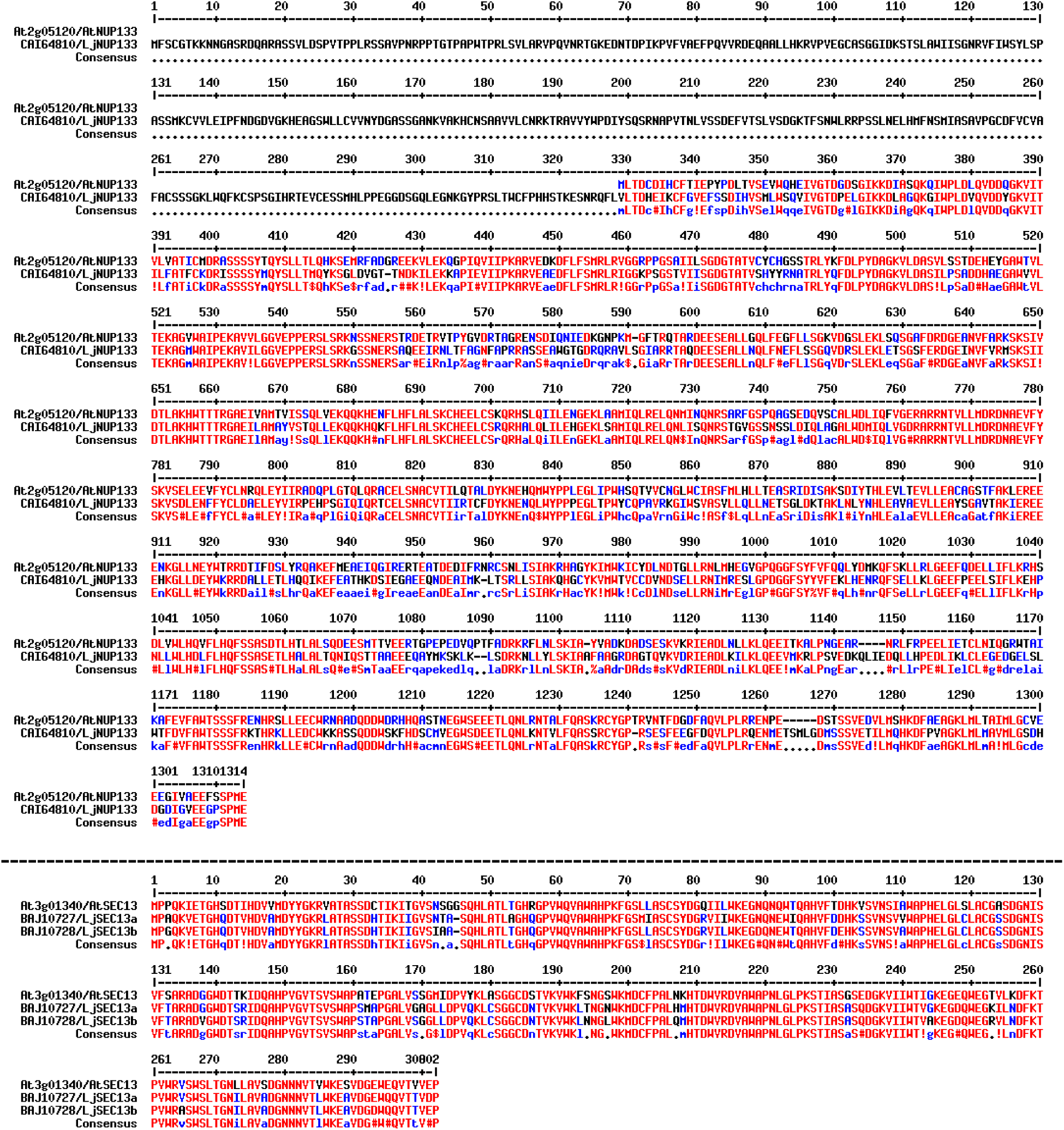
Alignments of the protein sequences of *L. japonicus* CSGs and closely related *A. thaliana* SNUPO genes. TAIR/GenBank protein identifiers are shown. Red, consensus level high = 90%; blue, consensus level low = 50%. Alignments were generated using the multiple sequence alignment software by Corpet (Corpet, 1988).

**S2 Fig.**
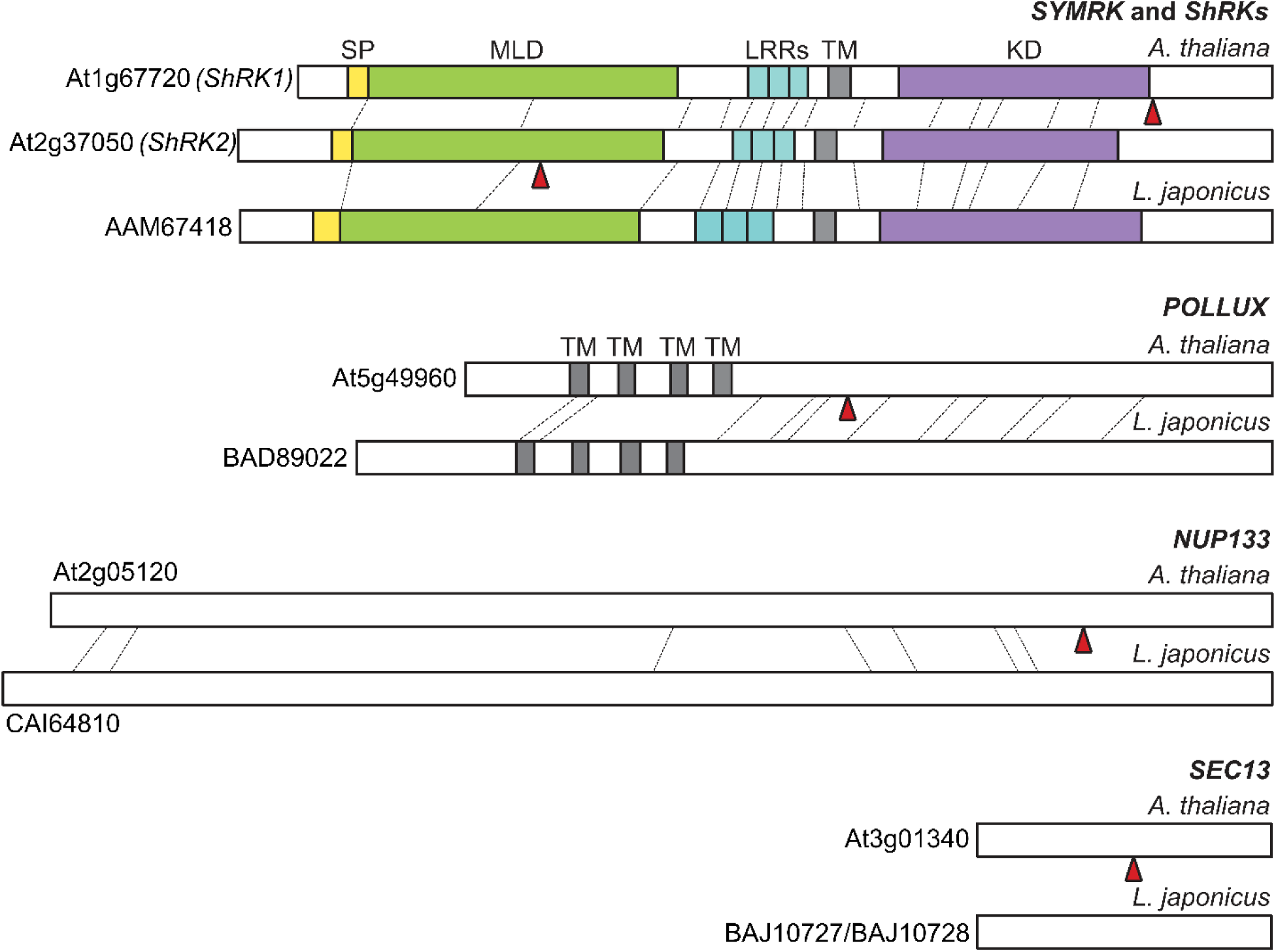
Comparison of gene structures and protein domains of *L. japonicus* CSGs and closely related *A. thaliana* SNUPO genes. TAIR/GenBank protein identifiers are shown; dashed lines, positions of introns in the original gene sequence; red triangles, positions of T-DNA insertion in the respective mutants; SP, signal peptide; MLD, malectin-like domain; LRRs, leucine-rich repeats; TM, transmembrane domain; KD, kinase domain.

**S3 Fig.**
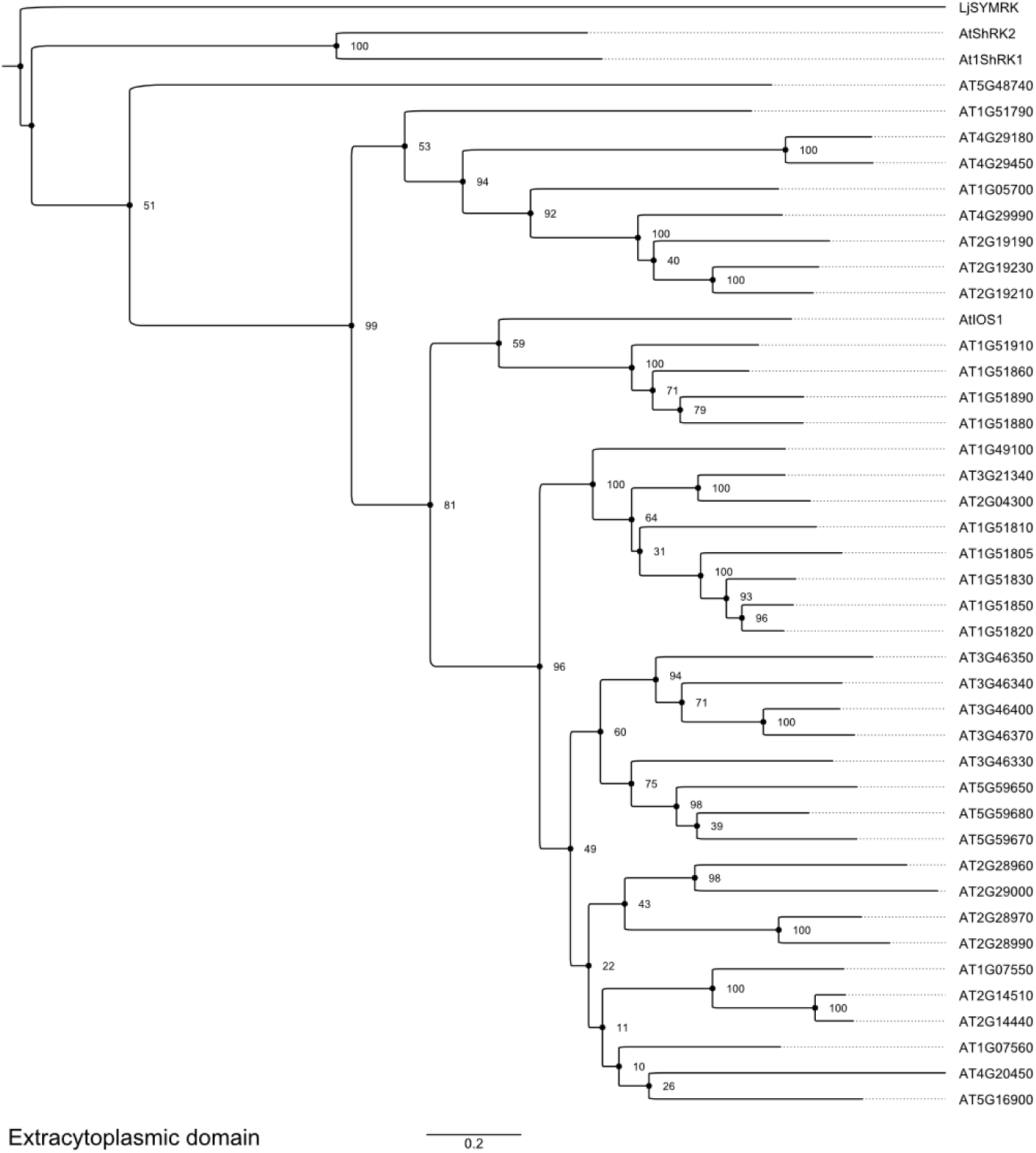
Maximum likelihood phylogenetic tree of the extracytoplasmic domains of MLD-LRR-RKs from *A. thaliana* and *L. japonicus* SYMRK. The specific domain composition featuring a MLD followed by the conserved gly-asp-pro-cys (GDPC) motif known to be required for the proper establishment of the symbiotic program in the root epidermis (Kosuta *et al.*, 2011; Antolín-Llovera *et al.*, 2014), and LRRs is not restricted to SYMRK but can also be found in 42 of the 50 members of LRR I-RLKs present in *A. thaliana* (Shiu & Bleecker, 2003; Hok *et al.*, 2011). A maximum likelihood phylogenetic tree based on the extracytoplasmic regions (excluding the signal peptides) of 42 MLD-LRR-RK proteins from *A. thaliana* and SYMRK from *L. japonicus*. Numbers on each node represent the respective bootstrap values. Bar, relative genetic distance (arbitrary unit).

**S4 Fig.**
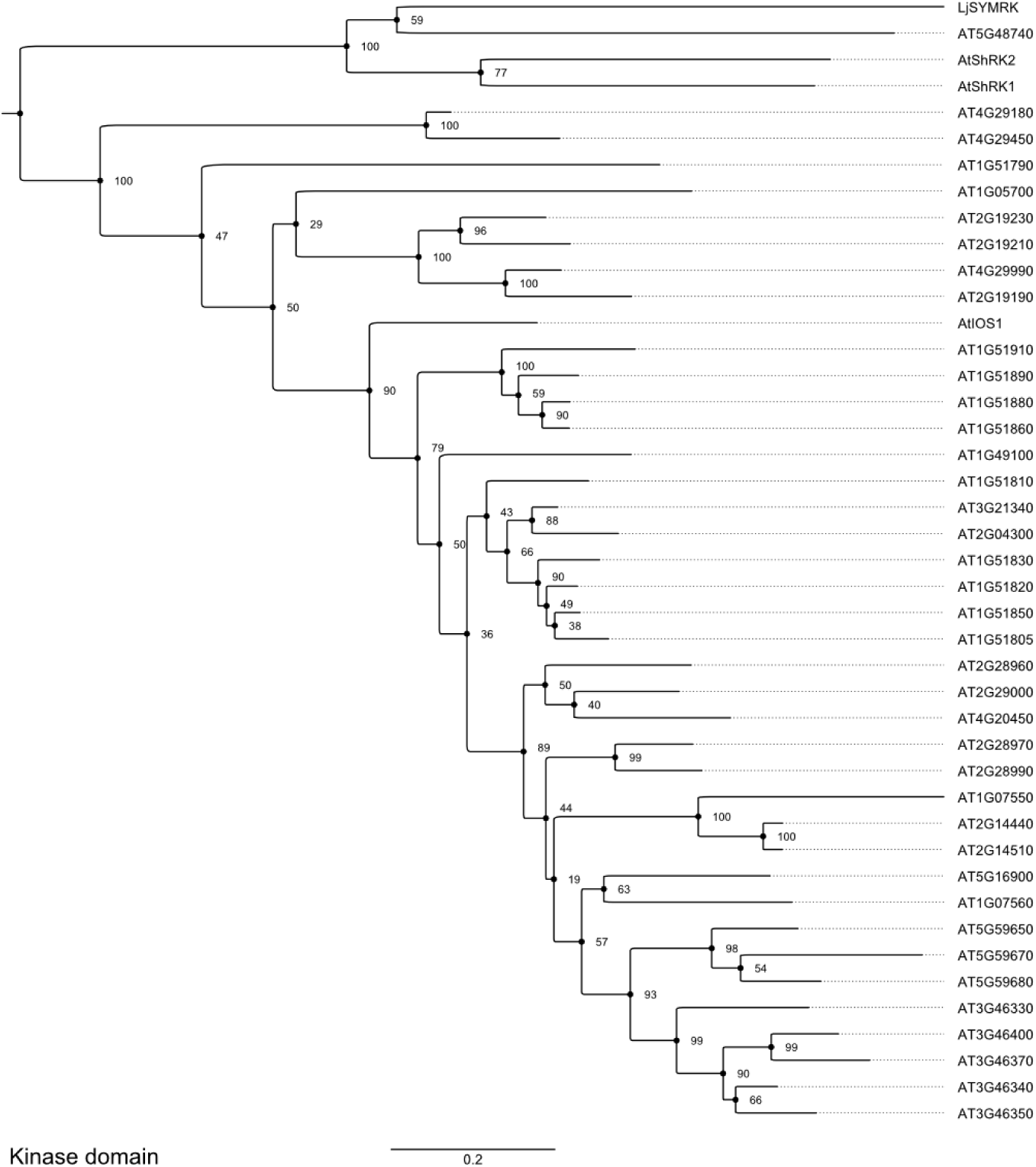
Maximum likelihood phylogenetic tree of the highly conserved kinase domains of MLD-LRR-RKs from *A. thaliana* and *L. japonicus* SYMRK. A maximum likelihood phylogenetic tree based on the highly conserved kinase domains of 42 MLD-LRR-RK proteins from *A. thaliana* and SYMRK from *L. japonicus*. Numbers on each node represent the respective bootstrap values. Bar, relative genetic distance (arbitrary unit).

**S5 Fig.**
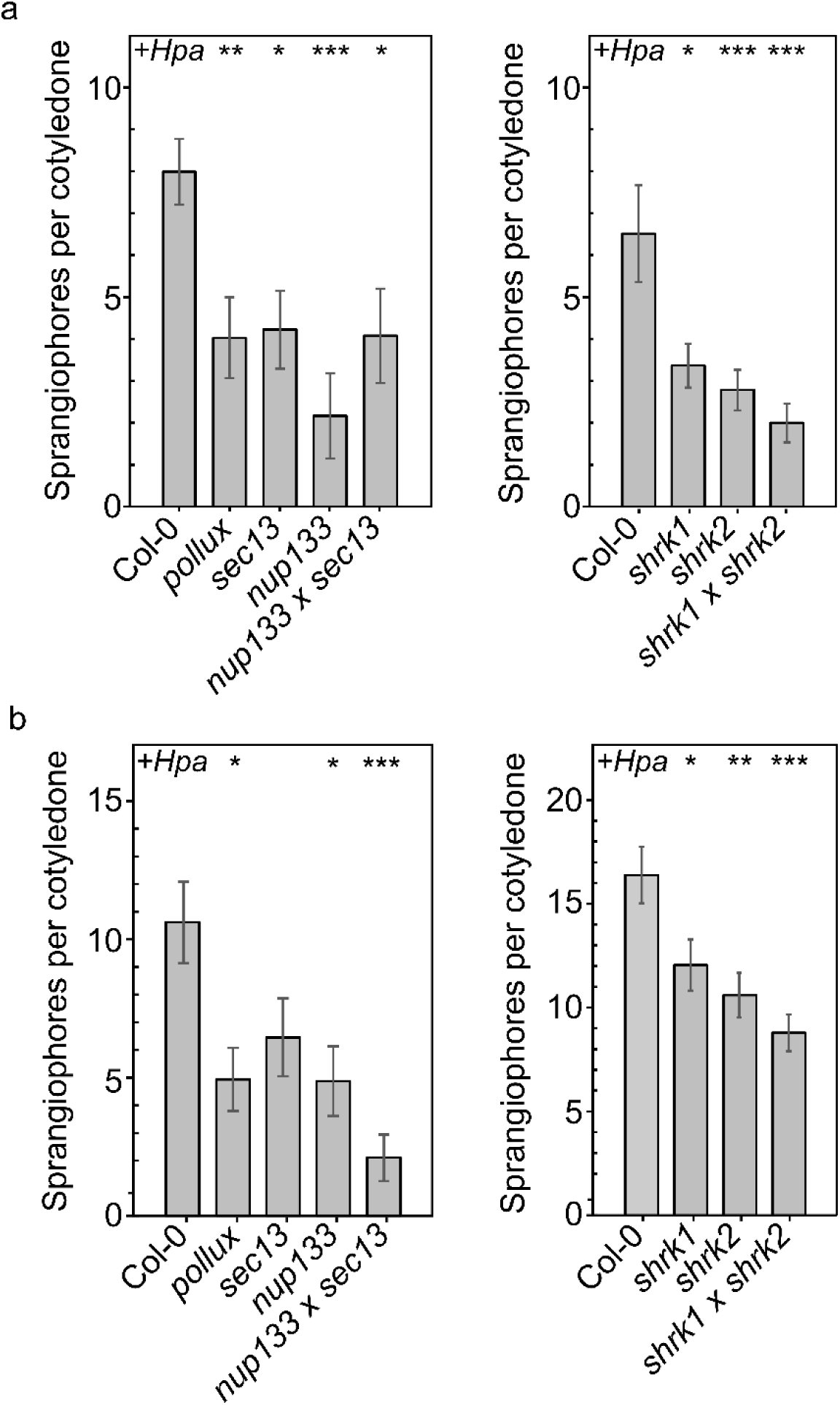
Mutations in A. thaliana SNUPO genes reduce the reproductive success of the oomycete downy mildew pathogen H. arabidopsidis. Bar charts represent the mean number of sporangiophores ± s.e.m on infected cotyledons of *A. thaliana* wild-type (Col-0) or the indicated mutants 4 dpi with *Hpa* isolate NoCo2 in two additional replicates (a + b). n = 21 – 99. Stars indicate significant differences to Col-0 (Wilcoxon– Mann–Whitney test with Bonferroni-Holm correction; *, p < 0.05; **, p < 0.01; ***, p < 0.001).

**S6 Fig.**
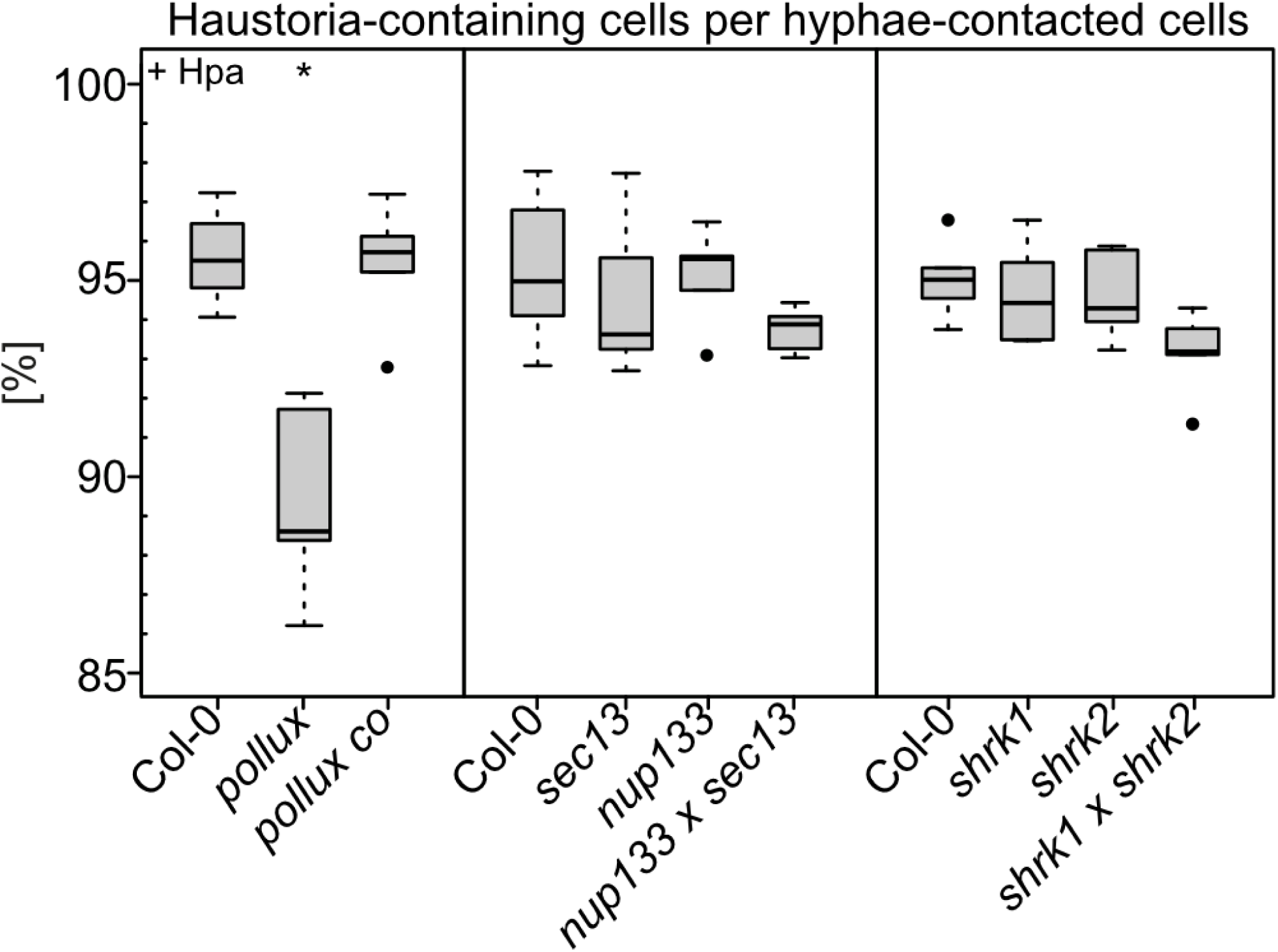
Mutations in *A. thaliana* SNUPO genes do not reduce the number of cells that accommodate haustoria. Boxplots represent the percentage of haustoria-containing cells per cells contacted by hyphae on leaves of *A. thaliana* wild-type (Col-0), the indicated mutants and the transgenic complementation lines (*pollux co, pPOLLUX:POLLUX*) 5 dpi with *Hpa* isolate NoCo2. For each genotype, at least ten independent stretches of hyphae per leaf have been analysed on at least 5 leaves. Black circles, data points outside 1.5 IQR of the upper/lower quartile; bold black line, median; box, IQR; whiskers, lowest/highest data point within 1.5 IQR of the lower/upper quartile. Stars indicate significant differences to Col-0 (Wilcoxon–Mann–Whitney test with Bonferroni-Holm correction; *, p < 0.05).

**S7 Fig.**
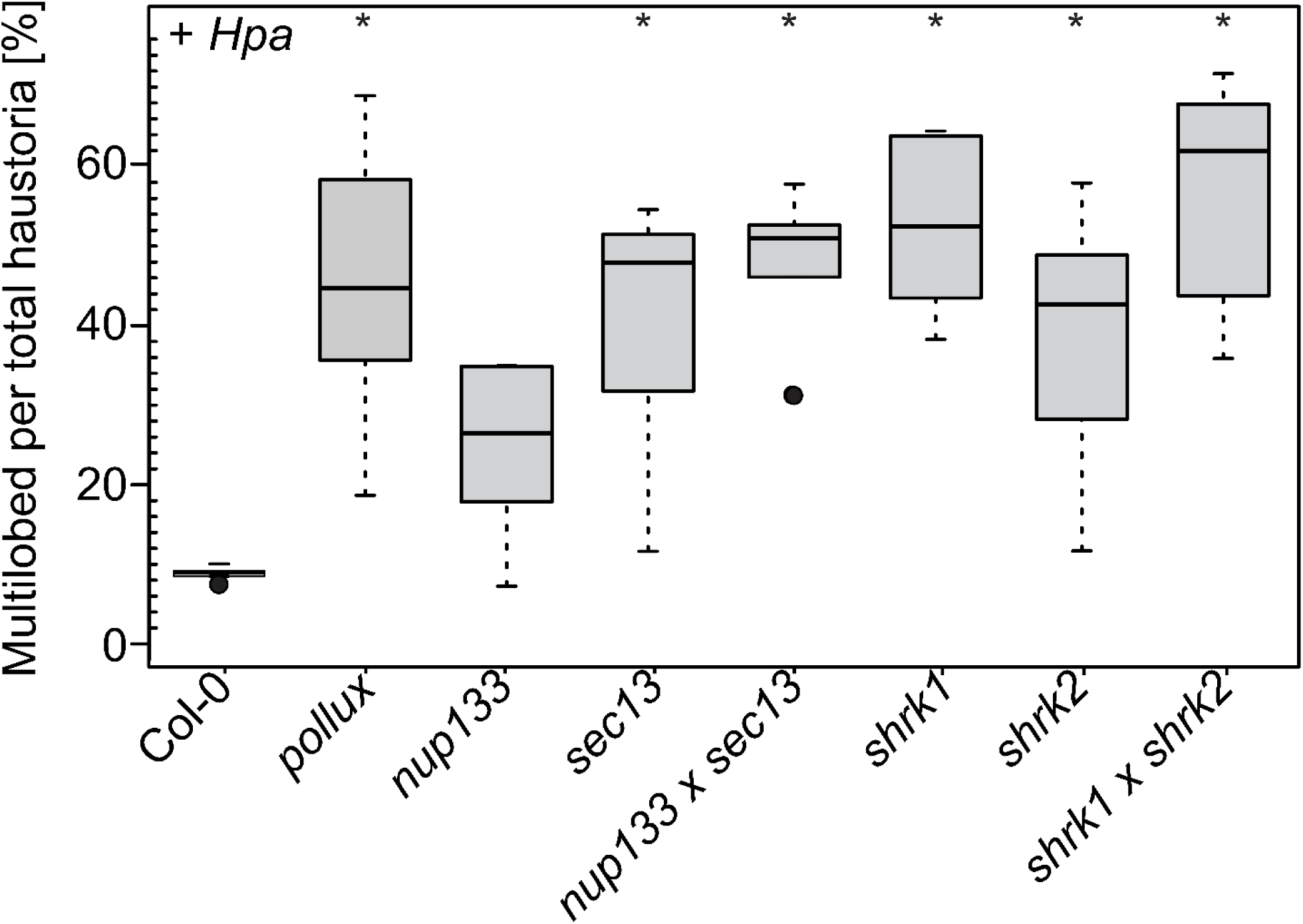
The morphology of *Hpa* haustoria is altered in *A. thaliana* SNUPO mutants. Boxplots represent the percentage of multilobed haustoria among total haustoria on *A. thaliana* wild-type (Col-0) or the indicated mutants 5 dpi with *Hpa* isolate NoCo2. For each genotype, at least ten independent stretches of hyphae per leaf have been analysed on at least 5 leaves. On average, 1100 haustoria have been analysed per genotype. Black circles, data points outside 1.5 IQR of the upper/lower quartile; bold black line, median; box, IQR; whiskers, lowest/highest data point within 1.5 IQR of the lower/upper quartile. Stars indicate significant differences to Col-0 (Wilcoxon–Mann–Whitney test with Bonferroni-Holm correction; *, p < 0.05).

**S8 Fig.**
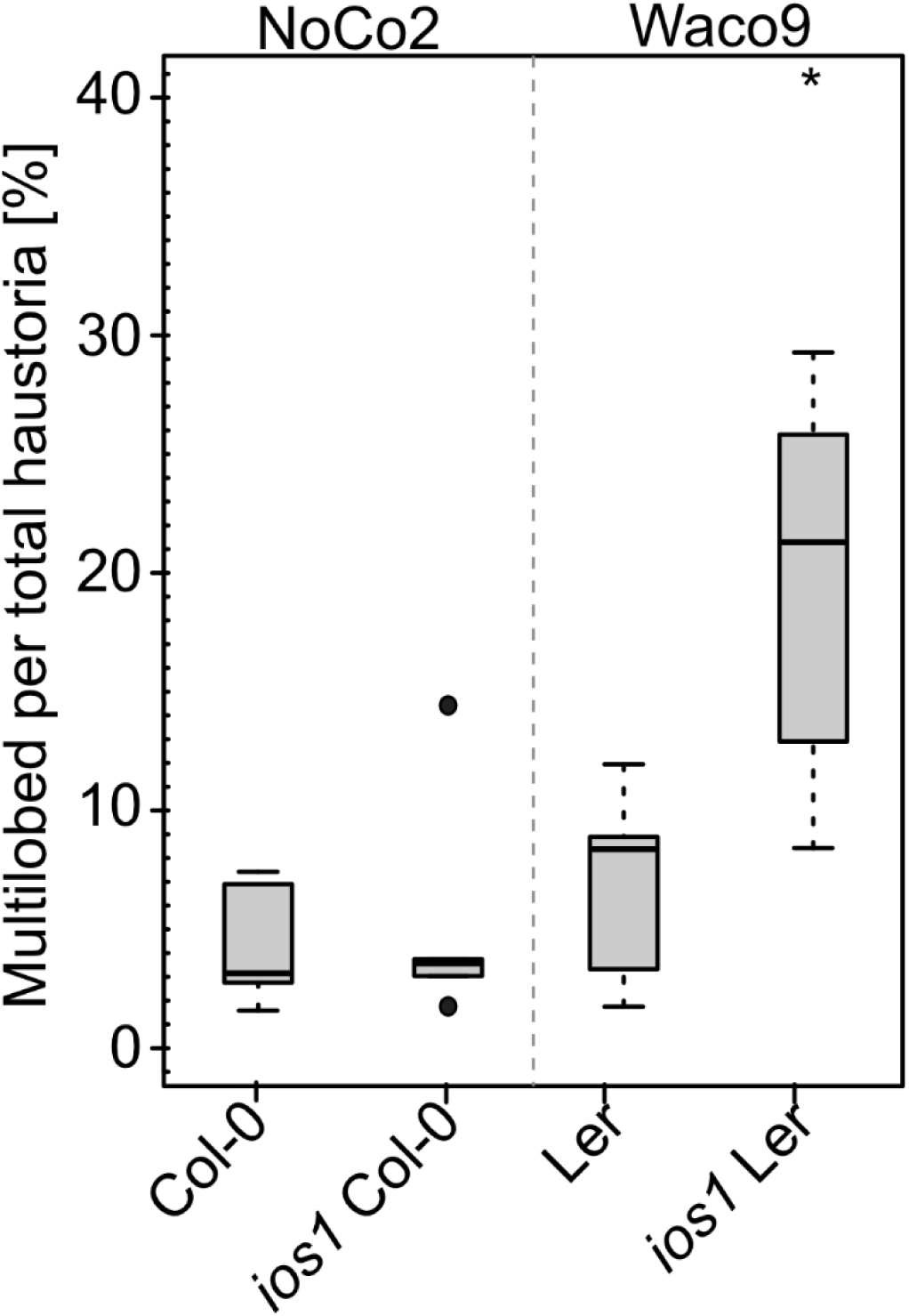
An increased frequency of multilobed *Hpa* haustoria in a *A. thaliana ios1* mutant was observed in the Ler background infected with Waco9, but not in the Col-0 background infected with NoCo2. Boxplots represent the percentage of multilobed haustoria among total haustoria on *A. thaliana* wild-type (Col-0; Ler), and *ios1* mutant lines 5 dpi with *Hpa* isolate NoCo2 (a), 5 dpi with *Hpa* isolate NoCo2 and Waco9. For each genotype, at least ten independent stretches of hyphae per leaf have been analysed on at least 5 leaves. Black circles, data points outside 1.5 IQR of the upper/lower quartile; bold black line, median; box, IQR; whiskers, lowest/highest data point within 1.5 IQR of the lower/upper quartile. Stars indicate significant differences to Col-0 (Wilcoxon– Mann–Whitney test with Bonferroni-Holm correction; *, p < 0.05).

**S9 Fig.**
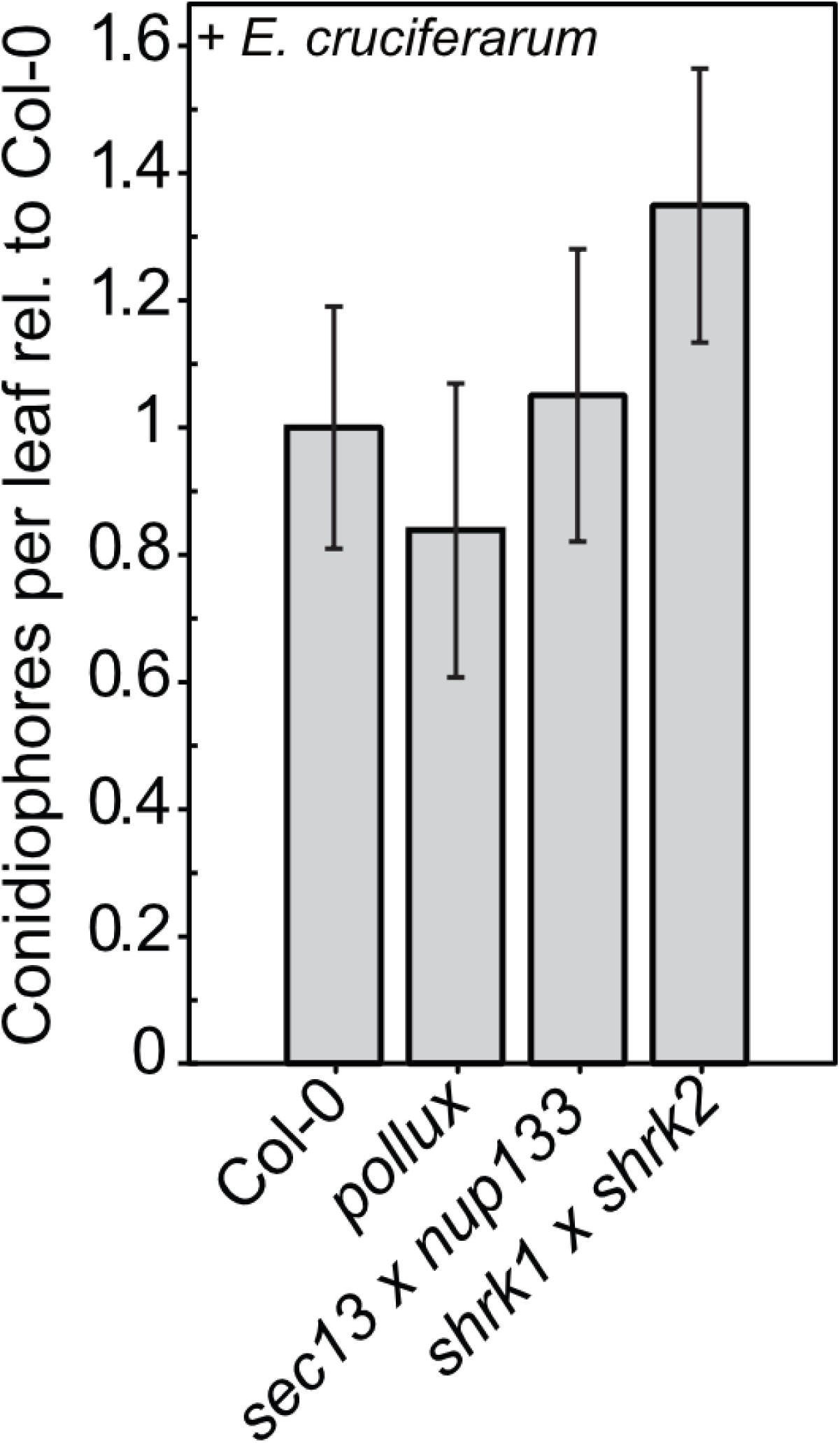
Mutations in *A. thaliana* SNUPO genes do not impair the reproductive success of the fungal powdery mildew pathogen *E. cruciferarum*. Bar charts represent the mean number of conidiophores ± s.e.m on leaves of the indicated mutants relative to the wild-type (Col-0) 5 dpi with *E. cruciferarum*. At least ten colonies per leaf on at least 6 leaves have been counted. No significant differences to Col-0 were detected.

**S10 Fig.**
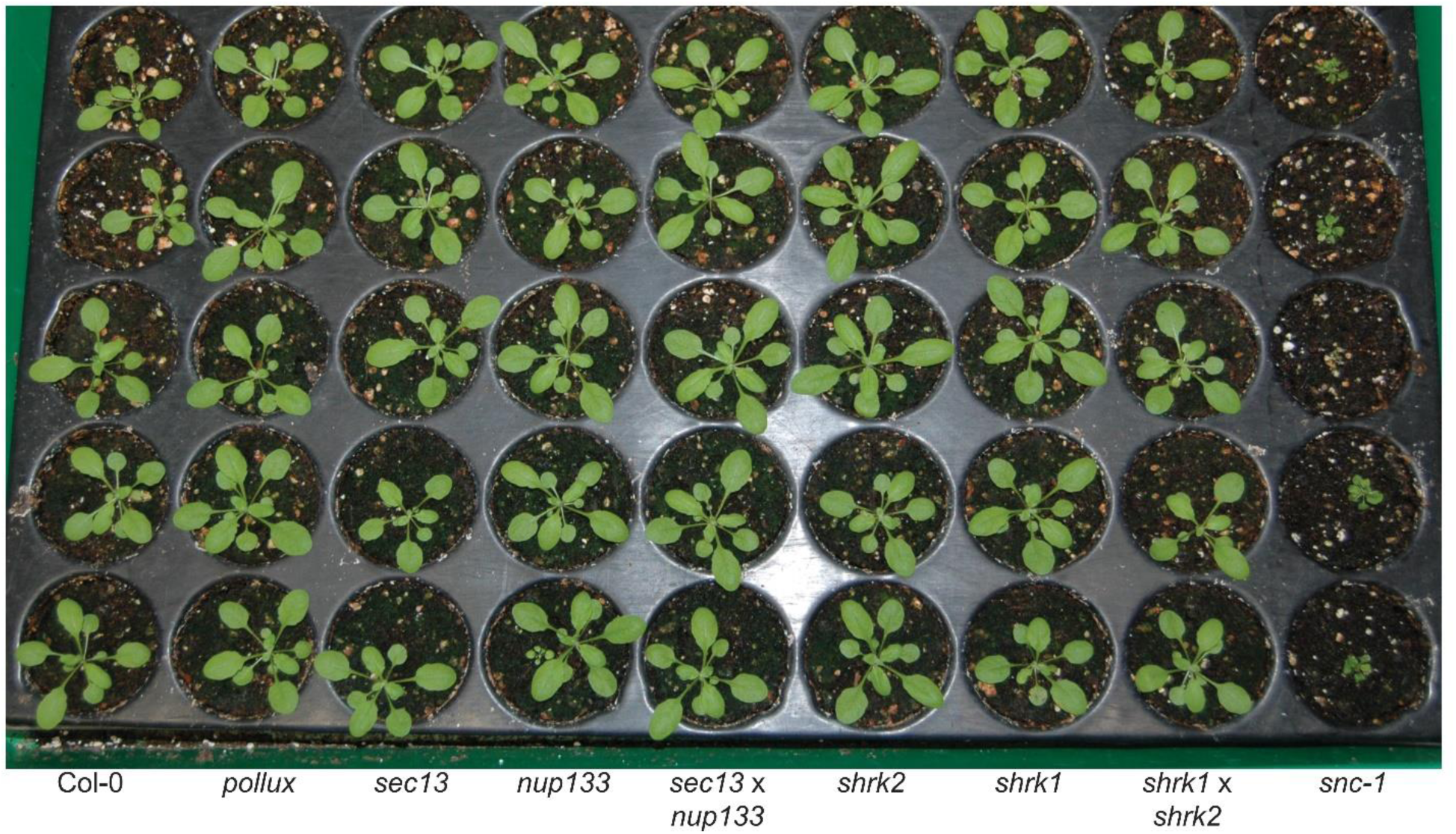
*A. thaliana* SNUPO mutants do not show developmental or growth defects. 3-week-old *A. thaliana* wild-type (Col-0) plants grown alongside the indicated mutant lines under long day conditions. The dwarf phenotype of *snc1* is included on the far right for comparison.

**S1 Table.**
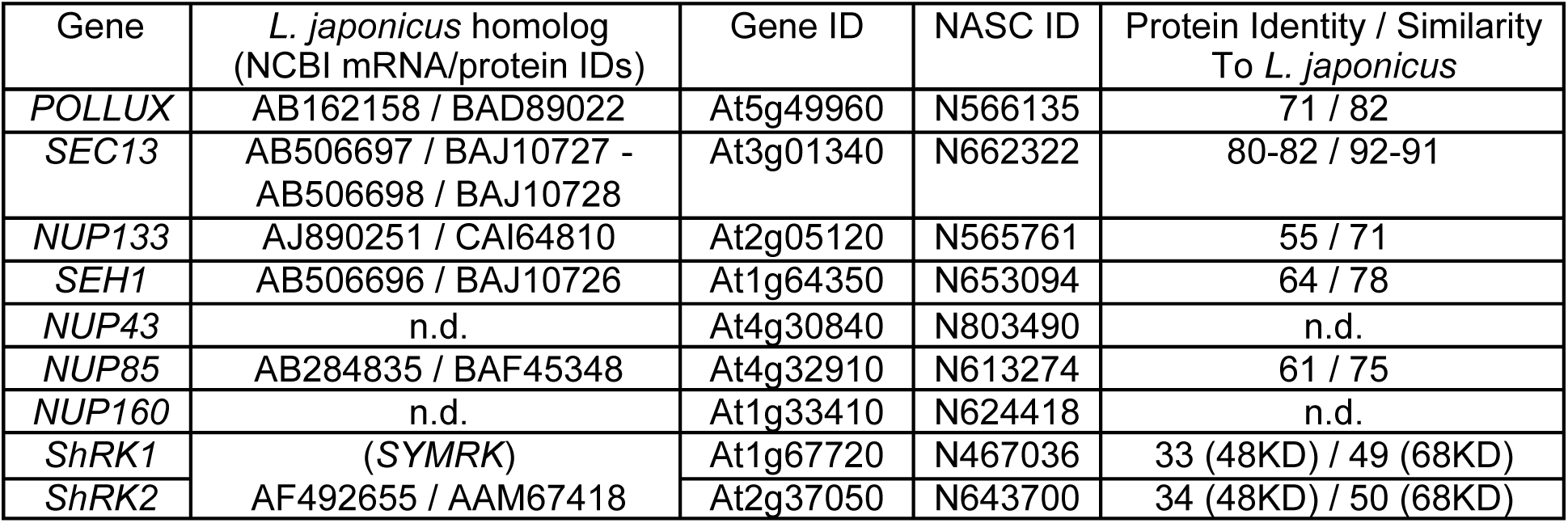
*A. thaliana* SNUPO genes with encoded proteins and their respective identities / similarities to related *L. japonicus* genes. Sequence identifiers and identities / similarities shared between the protein sequences of *A. thaliana* SNUPO and other nucleoporin genes and those of their respective *L. japonicus* counterparts. The numbers for AtSEC13 indicate the identity / similarity of its amino acid sequence to each of the two predicted LjSEC13 proteins. For ShRK1 and ShRK2, numbers in brackets refer to their kinase domain (KD) only. n.d., not detected. NASC ID: insertion mutant identifier.

**S2 Table.**
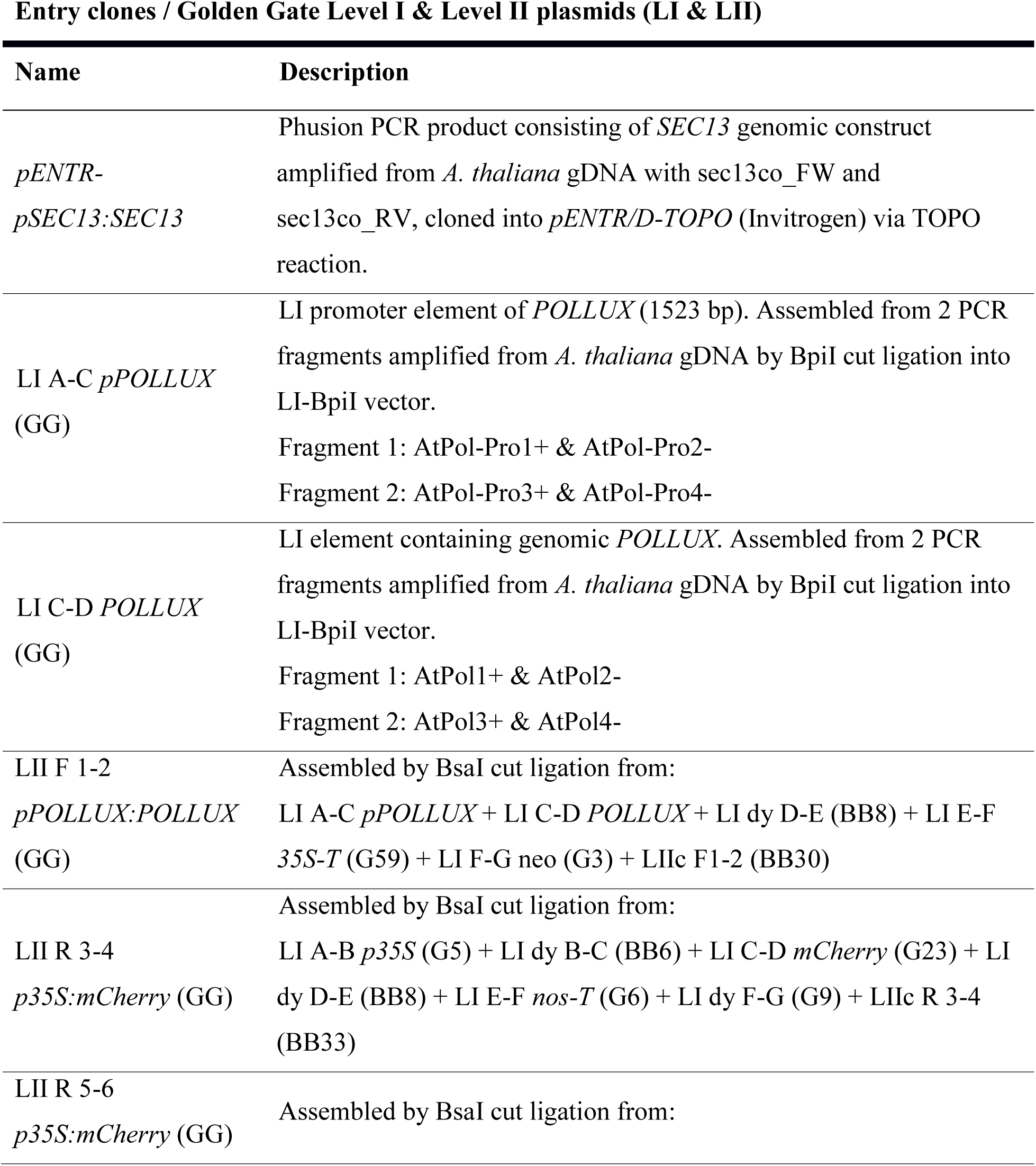
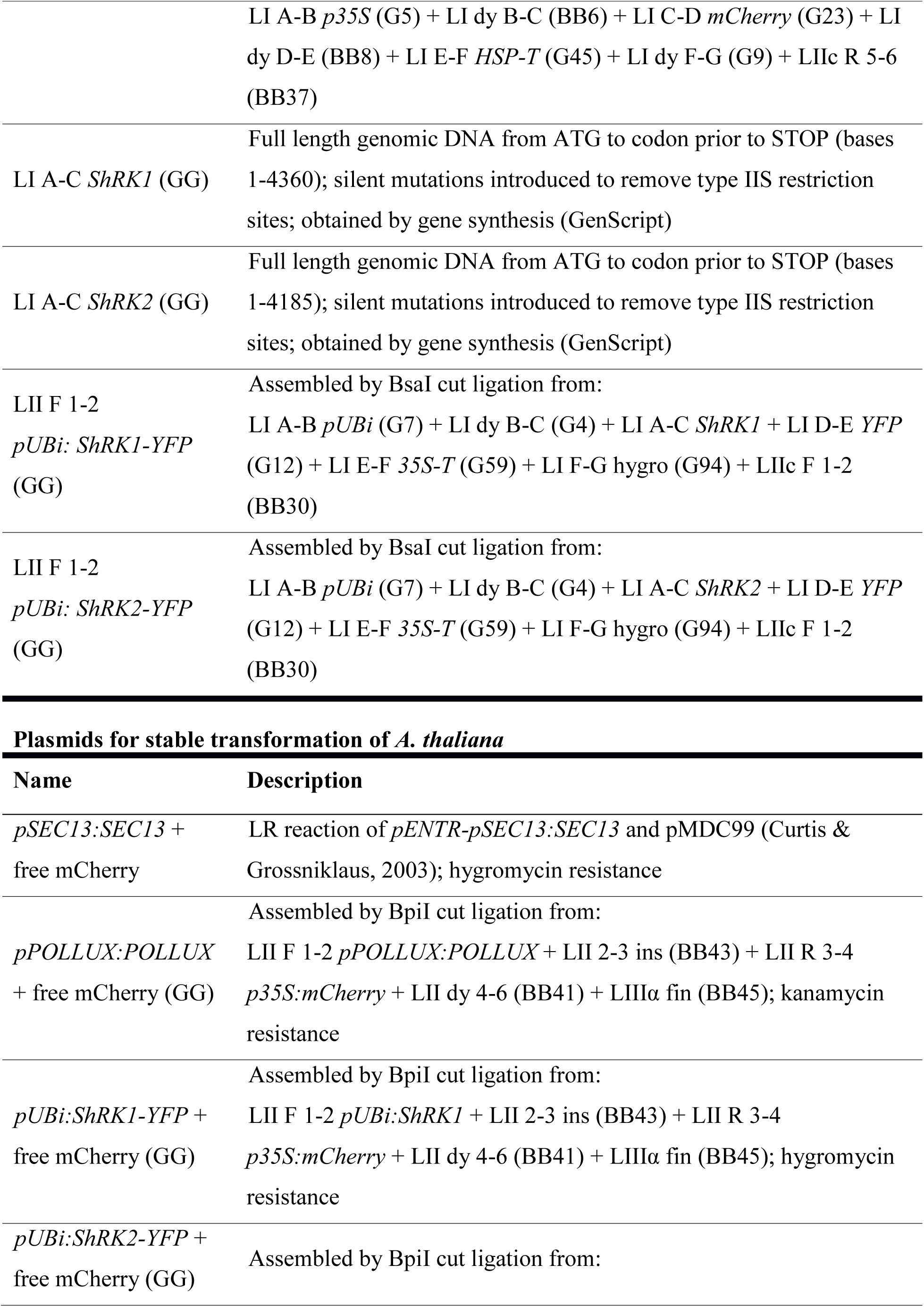
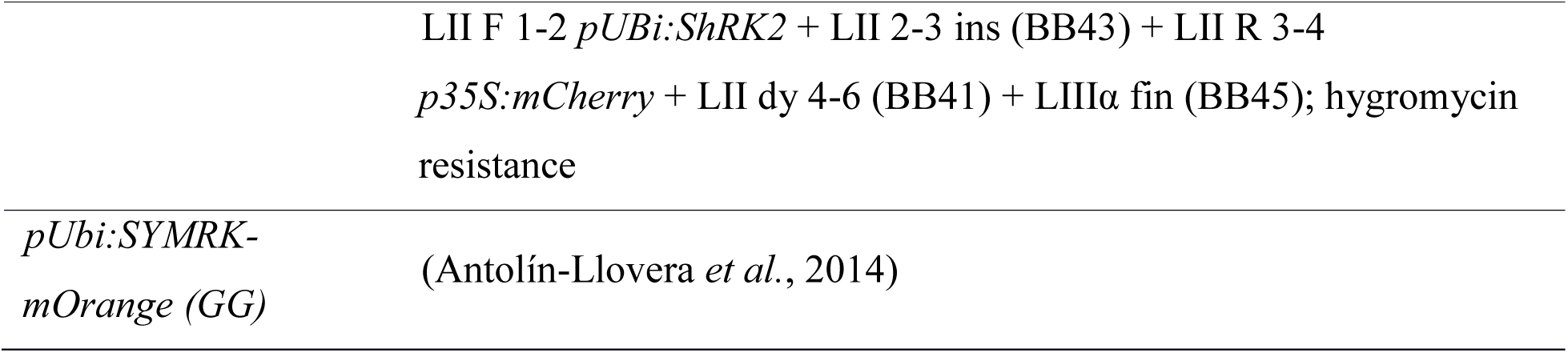
Constructs and cloning strategy. Constructs labelled with “GG” were generated via Golden Gate cloning. For details on assembly method, general modules and plasmids (Gxx, BBxx), see (Binder *et al.*, 2014). Golden Gate constructs contain silent mutations to facilitate cloning.

**S3 Table.**
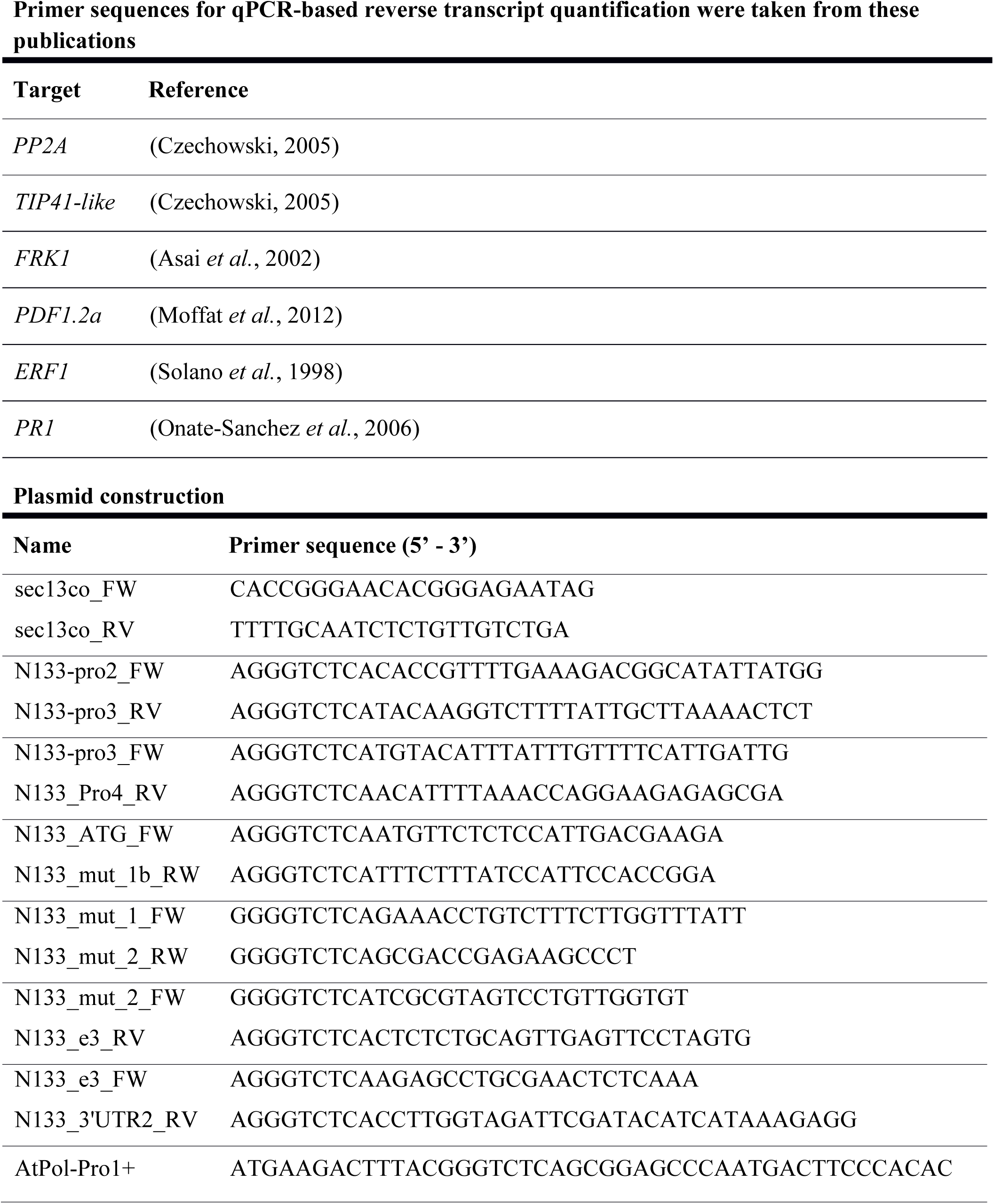
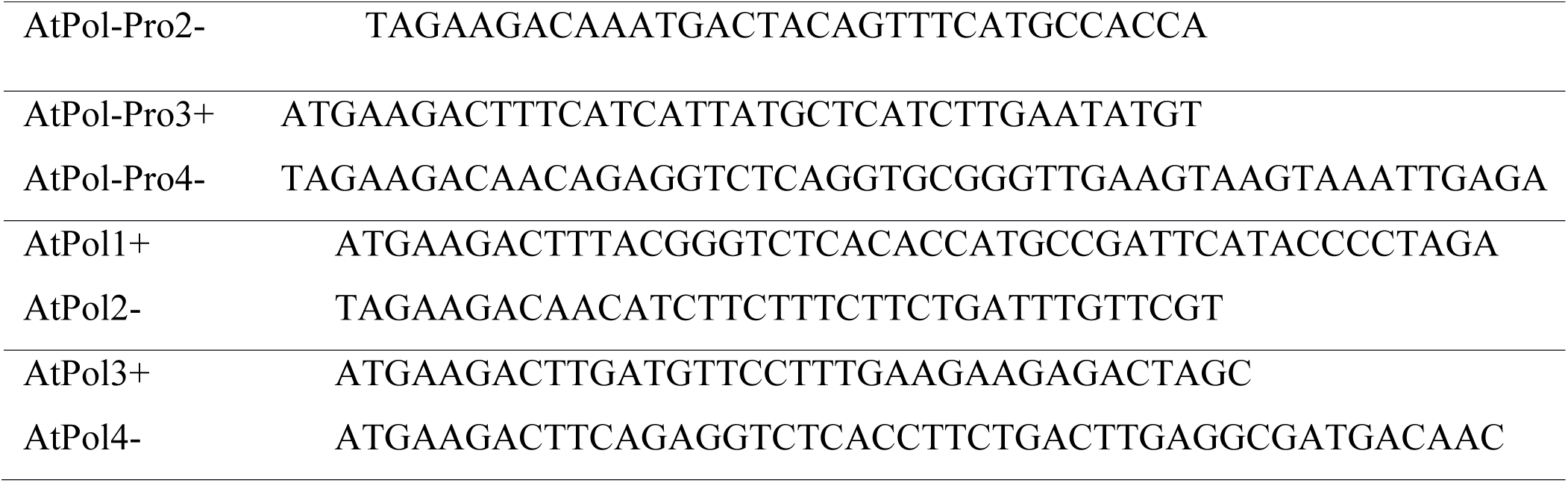
Oligonucleotides.

**S4 Table.** Mutant lines of *A. thaliana* SNUPO genes and seeds used in the study.

| Gene and ID | Mutant line | Seedbag |
| --- | --- | --- |
| *ShRK1* (At1g67720) | GK-699C04 2 | 1898 |
| *ShRK2* (At2g37050) | SALK_143700 | 1902 |
| *ShRK1* (At1g67720) x *ShRK2* (At2g37050) | GK-699C04 2 x SALK_143700 | 2118 |
| *Sec13* (At3g01340) x *Nup133* (At2g05120) | SALK_045825C x SALK_092608C | 2130 |
| *Nup133* (At2g05120) | SALK_092608C | 1941 |
| *Pollux* (At5g49960) | SALK_066135C | 2112 |
| *ShRK1* co (At1g67720) | GK-699C04 2 complemented with<br>*pUBi:ShRK1-YFP* + free mCherry | 2089 |
| *ShRK2* co (At2g37050) | SALK_143700 complemented with<br>*pUBi:ShRK2-YFP* + free mCherry | 2079 |
| *Sec13* co (At3g01340) | SALK_045825C complemented<br>with *pSEC13:SEC13* + free mCherry | 2026 |
| *Pollux* co (At5g49960) | SALK_066135C complemented<br>with *pPOLLUX:POLLUX* + free<br>mCherry | 2149 |

